# Super-resolution imaging flow cytometry

**DOI:** 10.64898/2026.10.18.759815

**Authors:** Yingdong Luo, Tianben Ding, Yuhong Liu, Yaqi Zhao, Hanqing Wang, Masako Nishikawa, Takamasa Ishino, Yosuke Togashi, T. Nicolai Siegel, Keisuke Goda

**Author notes:** Corresponding authors (T. Ding), (K. Goda).

## Abstract

Organelle architecture is a key determinant of cellular state. However, its heterogeneity remains difficult to quantify at population scale because conventional imaging flow cytometry is diffraction-limited. Here we present super-resolution imaging flow cytometry (SR-IFC), a high-throughput structured-illumination platform combining stroboscopic excitation, microfluidic hydrodynamic focusing and AI-assisted reconstruction. From three phase-shifted frames acquired along the flow axis, SR-IFC reconstructs images with approximately 118-nm isotropic in-plane resolution, about two-fold beyond the diffraction limit, at up to approximately 720 cells per second. In microbiology, SR-IFC resolves individual lipid droplets in nitrogen-starved *Euglena gracilis* and reveals spatial reorganization during lipid biogenesis missed by diffraction-limited imaging. In immunology, it maps activation-associated mitochondrial remodeling across primary human CD8^+^ T-cell differentiation, including a transient networked state. SR-IFC thus provides a broadly applicable platform for population-scale super-resolution phenotyping across diverse areas of biology.

## INTRODUCTION

A central theme across modern cell biology is that genetically identical cells can diverge widely in behavior and fate^1,2^, with much of this variability reflected at the level of their organelles^3,4^. Organelles are the functional compartments of the cell, and their abundance, morphology, and spatial arrangement set the metabolic, signaling, and biosynthetic capacity of each cell^5,6^. Cell-to-cell differences in these organelle properties, termed organelle heterogeneity, are increasingly recognized as drivers of biological functions, influencing how immune cells commit to memory or effector fates, how tumor cells adapt to stress and therapy, and how microbial and algal cells can be engineered to accumulate valuable products^7^. Many of the features that distinguish one cell from another are only tens to hundreds of nanometers in scale, below the diffraction limit of conventional microscopy. At the same time, biologically important states, such as rare subpopulations and transient intermediates, become apparent only when large numbers of cells are examined. Capturing organelle heterogeneity therefore requires two capabilities at once: resolving subcellular detail within individual cells and sampling enough cells to reconstruct the full distribution of cellular states.

These requirements have long been difficult to satisfy simultaneously. Spatial resolution and throughput occupy opposite ends of a fundamental trade-off in bioimaging: because the amount of optical information that can be collected per unit time is finite^8^, imaging systems can typically be optimized either to resolve fine details or to measure many cells rapidly, but rarely both. Super-resolution microscopy occupies the high-resolution end of this spectrum and enables direct visualization of organelles^9–11^. However, it was primarily developed for static, mounted samples imaged one field of view at a time. Extending super-resolution microscopy to large cell populations generally requires mechanical scanning of the sample or stage, which re-couples throughput to the speed and precision of the scan^12–15^. Flow cytometry occupies the opposite end of the spectrum. By streaming cells in single file past a fixed optical probe, imaging flow cytometry (IFC) can measure hundreds to thousands of events per second and has become a powerful platform for large-scale single-cell analysis^16–19^, with recent advances extending it to multimodal, volumetric, label-free Raman, and fluorescence-lifetime contrast^20–24^. However, the images acquired by IFC remain diffraction-limited, restricting most analysis to coarse, cell-scale features such as size, aspect ratio, and overall morphology, and leaving sub-200 nm organelle structures unresolved^20,25^. Super-resolution has been brought into microfluidic flow in a few forms: point-spread-function engineering localizes sparse emitters in flowing cells with nanometer precision but cannot reconstruct dense organelle architecture^26^, whereas image-scanning and structured-light-sheet approaches double the resolution of flowing cells but at rates of about one to ten cells per second, two orders of magnitude below flow-cytometric throughput^27,28^. A general platform that combines sub-diffraction, direction-independent lateral resolution with true flow-cytometric throughput has therefore remained unavailable.

In this pape, we address this long-standing speed-resolution trade-off with super-resolution imaging flow cytometry (SR-IFC), a platform that resolves organelles with sub-diffraction, direction-independent lateral resolution while cells flow at cytometric speed. Instead of relying on mechanical scanning, SR-IFC captures three phase-shifted structured-illumination frames of each cell during its brief transit and reconstructs them into a single high-resolution image. Structured illumination modulated along the flow axis provides a physical, measured resolution enhancement along that axis, while a self-supervised reconstruction framework, trained solely on the measured data without any external training set, extends this information to recover equal resolution along the transverse axis in the focal plane. Stroboscopic excitation freezes cell motion, and three-dimensional hydrodynamic focusing keeps each cell within the depth of field, enabling sharp frame acquisition and accurate registration in flow. Together, these elements achieve an in-plane resolution of approximately 118 nm, corresponding to an approximately two-fold improvement over diffraction-limited wide-field (WF) imaging, at throughputs of up to approximately 720 cells per second.

We demonstrate the broad utility of SR-IFC in two distinct areas of biology. The two applications were chosen to represent two common types of organelle measurement that require high spatial resolution: quantifying the number and spatial distribution of small, discrete organelles, and classifying the morphology of interconnected organelle networks. In microbiology, SR-IFC resolves and counts individual lipid droplets in nitrogen-starved *Euglena gracilis* and reveals coordinated spatial reorganization during lipid biogenesis, providing a quantitative readout relevant to metabolic-engineering screens. In immunology, SR-IFC captures activation-associated mitochondrial remodeling across thousands of primary human CD8^+^ T cells and identifies image-defined morphological states along differentiation trajectories, including a transient networked state not resolved by diffraction-limited imaging. By combining super-resolution imaging with population-scale throughput, SR-IFC extends imaging flow cytometry from cell-scale morphology to organelle-resolved phenotyping and provides a broadly applicable framework for studying subcellular heterogeneity across diverse biological systems.

## RESULTS

### System design and working principles of SR-IFC

Resolving organelles below the diffraction limit in microscopy can be achieved in several ways^9,29^, but the constraints of flow cytometry sharply narrow the options. Single-molecule localization methods such as photoactivated localization microscopy (PALM) and stochastic optical reconstruction microscopy (STORM) provide the finest resolution, but they reconstruct images from thousands of sequential frames of stochastically blinking fluorophores^30,31^. Point-scanning methods such as stimulated emission depletion (STED) microscopy achieve nanoscale detail by raster-scanning a shaped focus across the sample^32,33^. Both approaches require dwell times, photon budgets, or specialized photoswitchable labels that are incompatible with cells crossing the field of view within milliseconds. Structured-illumination microscopy (SIM) sits at a different position in this trade space: it is a wide-field, camera-based technique that doubles resolution using only a small number of patterned-illumination frames, works with conventional dyes at modest light doses, and supports the short exposures required for high-speed flow^34,35^. These properties make SIM a feasible foundation for super-resolution imaging in flow, and SR-IFC builds on this principle. The key obstacle, however, is that conventional SIM typically requires multiple grating orientations, usually nine frames in total, to achieve isotropic in-plane resolution. This would be a bottleneck for imaging a cell in transit. Overcoming this limitation by achieving SIM with a minimal number of frames is therefore central to the SR-IFC design. In addition, super-resolution imaging in flow imposes three requirements that static microscopy does not face, and these requirements dictate the system architecture. First, the resolution-encoding measurement must be completed within the brief window in which a cell crosses the field of view, precluding long sequential acquisitions. Second, every frame must remain sharp despite continuous motion, because motion blur would erase the high-spatial-frequency information responsible for the resolution gain. Third, successive frames of the same cell must be accurately registered even as the cell moves between exposures. SR-IFC addresses these requirements with three specialized modules: an optical module that encodes sub-diffraction detail through structured illumination, a microfluidic module that stabilizes each cell within the focal volume, and a computational module that reconstructs an isotropic in-plane image from a minimal set of frames (Fig. 1).

**Fig. 1.**
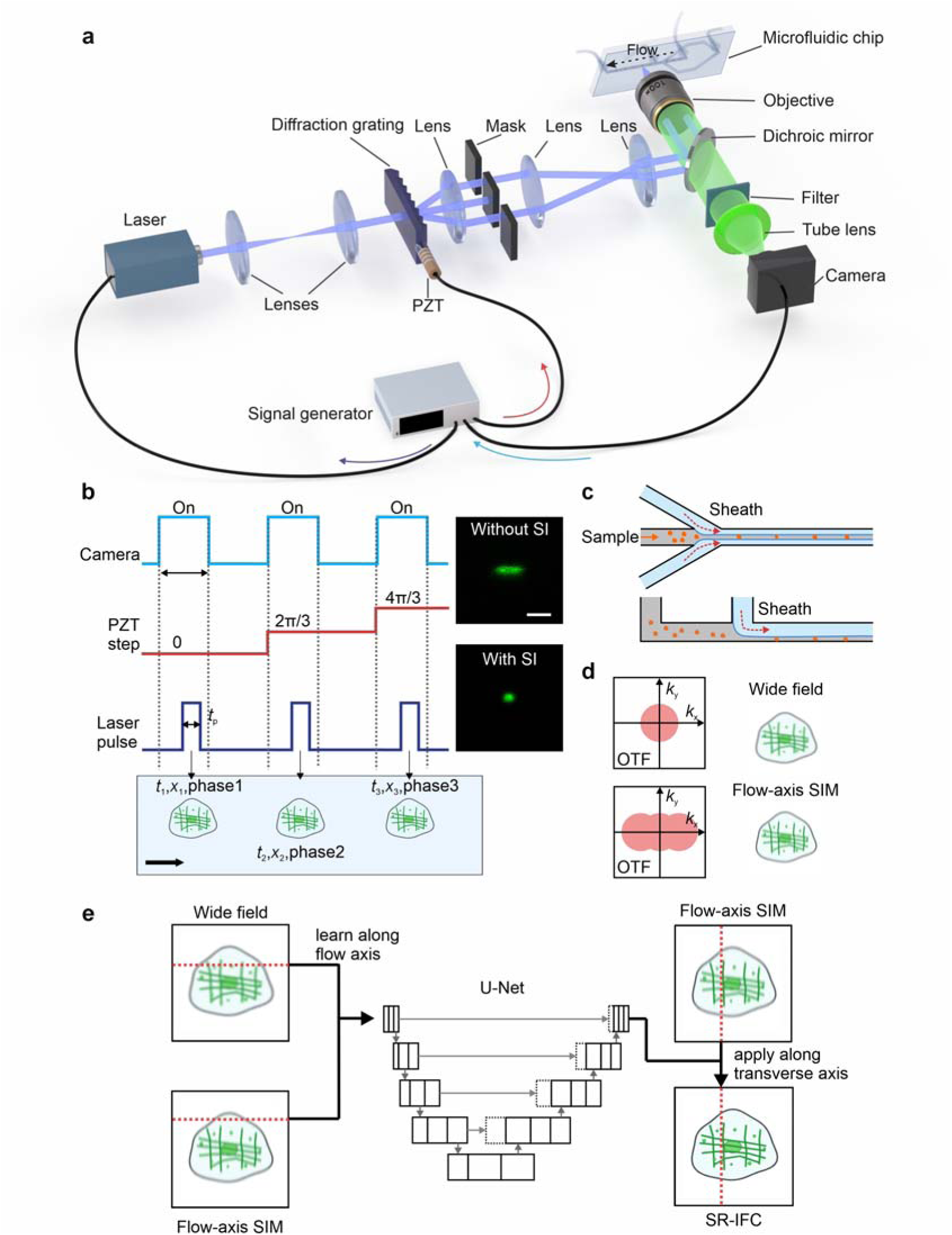
Working principles of SR-IFC. **a**, Optical system comprising a continuous-wave 488 nm laser, lenses, a diffraction grating mounted on a piezoelectric transducer (PZT), an order-selection mask, a microfluidic chip imaged through a 100× objective, a dichroic mirror, an emission filter, a tube lens, and a camera, with system synchronization controlled by a signal generator. **b**, Timing diagram of camera exposure, PZT-driven grating phase stepping, and 300 ns stroboscopic laser pulses used to acquire three phase-shifted frames. The cell advances along the flow axis between frames, such that each frame is recorded at a distinct cell position, acquisition time, and grating phase. Here, *t* = 300 ns denotes the laser-pulse width. Inset, 3 μm fluorescent beads flowing at 50 mm/s without and with stroboscopic illumination (SI), demonstrating motion-blur suppression. Scale bar, 5 μm. **c**, Three-dimensional hydrodynamic focusing of the sample stream by side and top sheath flows. **d**, Optical-transfer-function (OTF) support for WF imaging and flow-axis SIM, representing the range of spatial frequencies transmitted by each imaging mode. Flow-axis SIM extends the OTF support only along the flow axis (horizontal axis), thereby recovering finer spatial detail in that direction but not along the transverse axis. Right, schematic WF and flow-axis SIM images. **e**, Dual-input network reconstruction that combines the WF and flow-axis SIM images and recovers the missing transverse-axis detail in a single forward pass, producing an SR-IFC image with isotropic in-plane resolution. Center, simplified schematic of the U-Net-based network; the full architecture is shown in Supplementary Fig. 1.

The optical module is built on an inverted epi-fluorescence configuration with a 100×, 1.4-NA oil-immersion objective. A continuous-wave 488 nm laser is expanded and passed through a transmission grating mounted on a piezoelectric transducer (PZT), generating a pattern of illumination stripes whose intensity varies sinusoidally along the flow direction. Where the stripes overlap fine sample structures, they produce coarser moiré patterns that the objective can transmit; the reconstruction later recovers from these patterns detail finer than the diffraction limit. To remain within the cell-transit window, SR-IFC records only three phase-shifted frames at a single grating orientation, rather than the multiple orientations required for conventional SIM. Between consecutive camera frames, the PZT steps the grating so that the stripes advance along the flow axis by one-third of their period, while the cell moves along the same axis. Each of the three frames therefore captures the cell at a different position and with the stripes at a different position relative to the cell, that is, under a different illumination phase (Fig. 1b). In this way, the cell’s movement becomes part of the structured-illumination acquisition. Because continuous-wave excitation would smear the pattern at flow speed, each frame is exposed by a 300 ns stroboscopic laser pulse, which a common signal generator drives in synchrony with the camera and PZT. During this pulse, a cell traveling at 50 mm/s advances only approximately 15 nm, well below the 65 nm sample-plane pixel, so motion is effectively frozen and structured-illumination contrast is preserved^22^ (Fig. 1b). The microfluidic module confines the sample stream by three-dimensional hydrodynamic sheath focusing, keeping cells within the depth of field of the objective throughout the frame series so that all images remain in focus and can be registered^36^ (Fig. 1c). Together, these modules acquire three sharp, single-orientation structured-illumination frames of each cell during its millisecond-scale transit through continuous flow (see Methods, “Optical system and synchronization” and “Microfluidic device”).

Because the illumination is modulated along the flow axis, the three phase-shifted frames encode sub-diffraction information along the direction of cell motion, while the transverse axis remains diffraction-limited (Fig. 1d). Conventional SIM recovers that axis by acquiring additional grating orientations, which flow cannot accommodate. SR-IFC instead measures the flow axis and infers the missing transverse axis information through a two-step reconstruction (Fig. 1e). In the first step, a standard single-orientation SIM reconstruction^34,37–39^ returns the high spatial frequencies encoded along the flow axis to their true positions in the image, super-resolving that axis. We refer to this intermediate image as flow-axis SIM. In the second step, a U-Net-based convolutional network^40–42^ estimates the unmeasured transverse-axis detail. The network takes the WF and flow-axis SIM images as input, combines a directional branch aligned with the modulation (flow) axis with a two-dimensional branch spanning the entire image, and returns an image with matched, isotropic in-plane super-resolution in a single forward pass (Fig. 1e; Supplementary Fig. 1). A key property of this reconstruction is that it uses no external training data. The network is trained only on the WF and flow-axis SIM measurements of the cells being imaged, so the recovered transverse-axis detail reflects the structural statistics of the sample itself rather than a prior transferred from an unrelated dataset. Accordingly, the estimate is most reliable for structures well represented in the sample and most conservative for the features oriented purely along the unmeasured transverse axis. Crucially, this computational step adds no additional excitation orientations or optical hardware. The network architecture, training-set composition, and convergence behavior are reported in Methods (“AI-assisted isotropic reconstruction”) and Supplementary Fig. 1. The frame registration and the single-orientation SIM reconstruction are detailed in Supplementary Notes 1 and 2. The result is isotropic in-plane super-resolution acquired within a single cell’s transit, establishing the capability on which the remainder of this work builds.

### Validation across multiple simulation scenarios of SR-IFC

Because SR-IFC directly measures super-resolution information only along the flow axis, two questions must be addressed before applying it to cells: whether the flow-axis measurement recovers the same structural detail as conventional SIM along that axis, and whether the network accurately reconstructs the unmeasured transverse-axis information. Answering both questions requires images with exactly known ground-truth structures, which can only be obtained through simulation. We therefore generated fluorescent test objects, referred to as phantoms, and imaged them using a numerical model of the microscope, allowing each reconstruction to be compared directly with the known structure. The model reproduced the major steps involved in forming each raw image: excitation by the structured illumination pattern, blurring by the point-spread function, sampling by the camera pixels, and the addition of photon shot noise and camera read noise (Methods, “Simulation and image-quality metrics”; Supplementary Table 1). The phantoms were kept stationary so that reconstruction performance could be evaluated independently of cell motion, which is examined experimentally in the next section. From the same simulated raw frames, we generated four image modes and compared each with the noiseless ground truth, defined as the phantom projected onto the image plane without optical blur or noise (projected GT in Fig. 2a). WF imaging, defined as the average of the three raw frames, which cancels the illumination pattern, served as the diffraction-limited negative control. Conventional SIM, which acquires three illumination orientations with three phases each, for a total of nine raw frames, and applies a standard reconstruction, served as the positive control defining the target performance. Between these controls were two single-orientation, three-frame modes: flow-axis SIM, the physical reconstruction that super-resolves only the flow axis, and SR-IFC, which applies the network to the flow-axis SIM result to infer the transverse-axis information. Thus, SR-IFC aims to achieve the performance of nine-frame SIM using only a three-frame acquisition. To evaluate reconstruction performance along independent axes of difficulty, we designed three structurally distinct phantoms (Fig. 2a). The bead pair phantom (P1) is a 5 by 5 array in which each site contains two 100 nm beads. The surface-to-surface gap increases from 60 to 250 nm across rows, and the pair orientation rotates from the flow axis to the transverse axis across columns, allowing two-point resolution to be evaluated as a joint function of separation and direction. The ellipsoids (P2) span aspect ratios from 1:10, 1:5, 1:3 to 1:1 at in-plane orientations from 0 to 90 degrees, with an approximately 100 nm short axis, testing the recovery of anisotropic morphology. The frequency-target array (P3) contains line pairs with periods from 240 to 600 nm at orientations from 135 to 0 degrees, testing how faithfully fine periodic detail is restored. These phantoms follow test-object designs commonly used to benchmark microscope resolution, including point pairs, elongated objects, and line gratings with graded spatial periods. They also mimic structures relevant to organelle imaging, such as closely spaced puncta representing vesicles or lipid droplets, elongated organelles such as mitochondria, and regularly spaced fine features whose resolvability depends on spatial period.

**Fig. 2.**
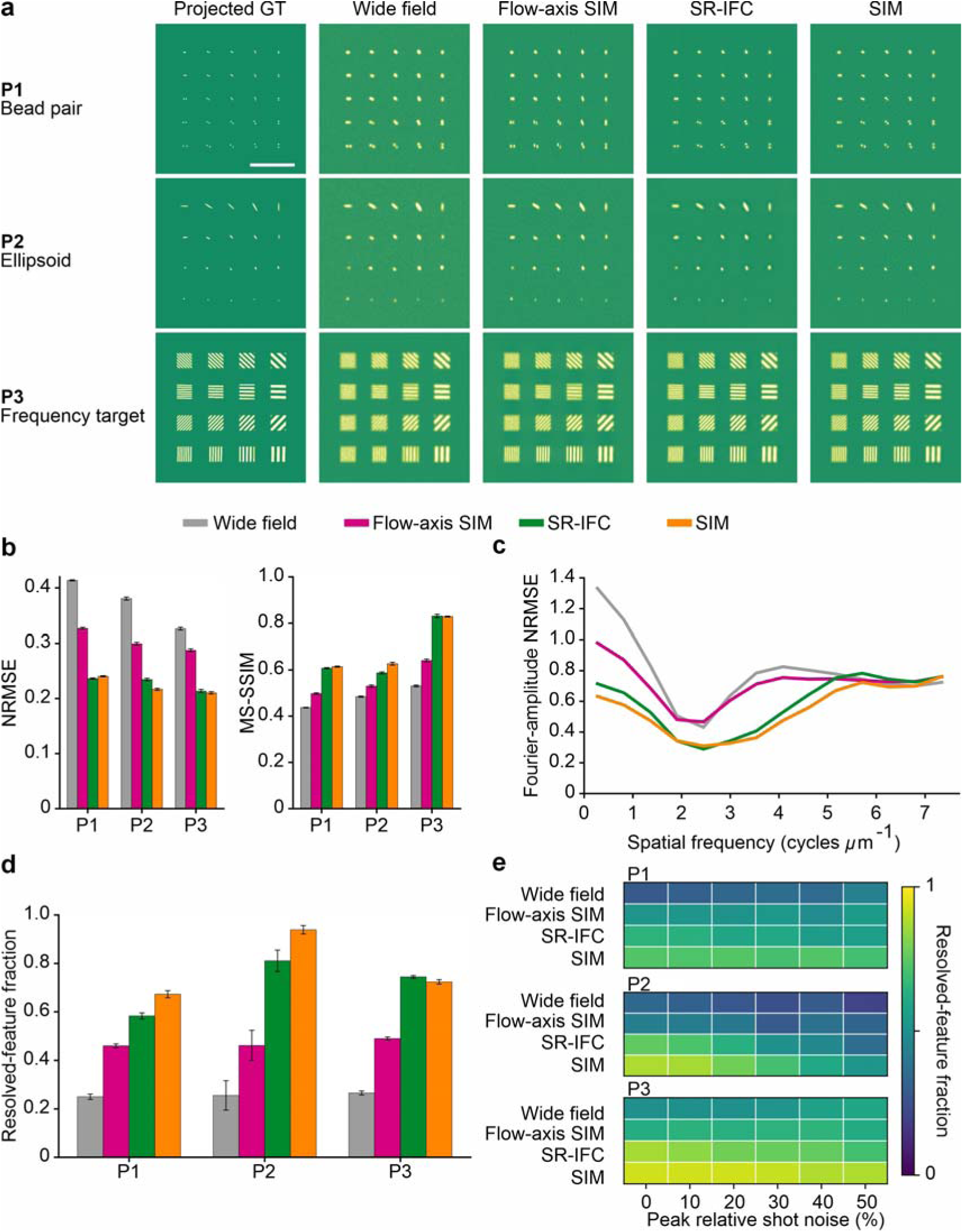
Validation across multiple simulation scenarios of SR-IFC. **a**, Bead pair (P1), ellipsoid (P2), and frequency target (P3) phantoms, each shown over the full field of view as the noiseless ground truth projected onto the image plane (projected GT) and as reconstructed by WF imaging, flow-axis SIM, SR-IFC, and nine-frame SIM. Scale bar, 5 μm. **b**, Pixel-level and structural fidelity against the ground truth, reported as normalized root-mean-square error (NRMSE) and multi-scale structural similarity (MS-SSIM) for each phantom and reconstruction mode. **c**, Fourier-amplitude NRMSE relative to the ground truth as a function of spatial frequency, averaged across the three phantoms. **d**, Task-specific resolution recovery for each phantom, reported as the resolved feature fraction, that is, the fraction of ground-truth features that meet the phantom-specific resolution criterion (see Methods, “Quantification of the resolved feature fraction”). **e**, Resolved feature fraction for each phantom and reconstruction mode as the peak relative shot noise increases from 0% to 50%, testing robustness to noise. In b and d, bars represent the mean ± standard error of the mean of 12 independent simulations.

SR-IFC reconstructions across the full field of view closely resembled conventional SIM, whereas the WF image was markedly blurred and flow-axis SIM sharpened structures along only one axis (Fig. 2a). We quantified this agreement using four families of metrics across all three phantoms, each obtained by comparing a reconstruction against its noiseless ground-truth image: pixel-level and structural fidelity in the object domain, spectral fidelity in the Fourier domain, a task-based resolution score, and robustness to added noise. In the object domain, we used two complementary metrics. The normalized root-mean-square error (NRMSE)^43^, for which lower values indicate better agreement, measures how accurately pixel intensities are reproduced and is therefore relevant to intensity- and size-based readouts such as organelle area. The multi-scale structural similarity (MS-SSIM)^44^, for which higher values indicate better agreement, measures how well edges, textures, and shapes are preserved across multiple spatial scales and therefore relevant to morphology-based readouts (both metrics are defined in Methods, “Simulation and image-quality metrics”). By both metrics, SR-IFC consistently outperformed flow-axis SIM and approached the performance of nine-frame SIM, with the largest remaining difference observed for the ellipsoid phantom (Fig. 2b). In the Fourier domain^45^, we compared the Fourier amplitude spectrum of each reconstruction with that of the ground truth as a function of spatial frequency. Because spatial frequency corresponds to the scale of image detail, a smaller amplitude error indicates more accurate recovery of information at that scale. Across the low-to mid-spatial-frequency range, the SR-IFC amplitude-error curve closely followed that of nine-frame SIM rather than those of WF or flow-axis SIM (Fig. 2c), indicating recovery of transverse spatial-frequency information missing from the latter two reconstructions. As a task-based measure of resolution^45^, we used the resolved feature fraction, defined as the proportion of known features, such as the bead pairs in P1 and the line pairs in P3, that each method resolves. The metric ranges from 0 to 1, with higher values indicating better resolution (Methods, “Quantification of the resolved feature fraction”). The same overall trend was observed. SR-IFC reached the SIM level for P1 and P2, slightly exceeded it for P3 (0.75 versus 0.72) and remained well above the WF floor for every phantom (Fig. 2d). Finally, because high-throughput flow imaging is inherently photon-limited, we reduced the simulated signal to assess noise robustness^46^. Specifically, we recalculated the resolved-feature fraction while increasing the peak relative shot noise from 0% to 50% (Methods, “Simulation and image-quality metrics”). The resolved feature fraction obtained with SR-IFC closely tracked that of the SIM reference across this noise range, indicating that the reconstruction remains stable under the low-light conditions in continuous flow (Fig. 2e). Zoomed regions of interest and orthogonal line profiles for each image mode are compared in Supplementary Fig. 2. Together, the object-domain, Fourier-domain, task-resolution and noise-robustness analyses across three structurally distinct phantoms show that SR-IFC closely approached the performance of conventional multi-orientation SIM. These results establish the central premise of the method: combining single-orientation physical SIM with a learned transverse-axis reconstruction is sufficient to recover isotropic in-plane super-resolution from only three frames across structurally diverse samples.

### Characterization of SR-IFC performance

We next experimentally quantified the spatial resolution achieved by SR-IFC. Individual 100 nm fluorescent beads were translated through the illumination field using a motion stage at approximately 50 mm/s, matching or exceeding the transit speed of cells in flow. Three consecutive phase frames were captured for moving beads (Fig. 3a), and the pipeline reconstructed WF, flow-axis SIM, and SR-IFC images, representing the diffraction-limited, single-orientation super-resolved, and isotropic in-plane super-resolved outputs, respectively (Fig. 3b). From the single-bead images, we extracted population-averaged point-spread functions and fitted them with two-dimensional Gaussians to obtain the full width at half maximum (FWHM) along each axis (Fig. 3c). SR-IFC achieved FWHMs of approximately 155 nm along the flow axis and 149 nm along the transverse axis, corresponding to spatial resolutions of approximately 118 nm and 110 nm, respectively, after deconvolution of the 100 nm bead diameter. In comparison, WF imaging of the same beads yielded resolutions of 222 to 227 nm, indicating an approximately twofold improvement (Fig. 3d). Two aspects of this result are particularly important. First, the flow- and transverse-axis resolutions differed by less than 10 nm, confirming that the learned transverse-axis reconstruction restores near-isotropic in-plane resolution on experimental data, not only in simulation. Second, this twofold improvement over WF imaging is consistent with the linear-SIM resolution limit, indicating that SR-IFC achieves from three single-orientation frames the resolution that conventional SIM typically requires nine frames to obtain.

**Fig. 3.**
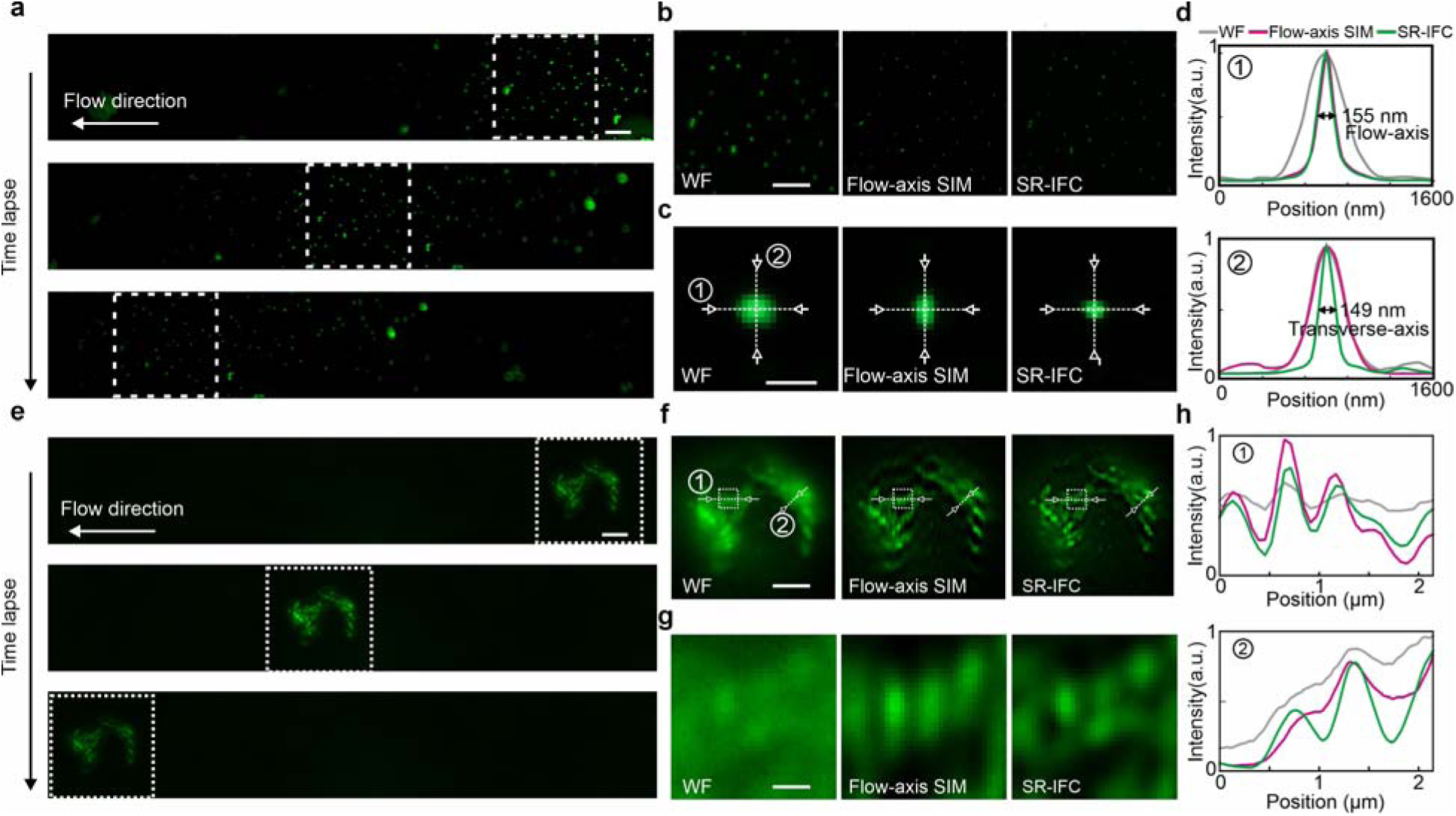
Characterization of SR-IFC performance. **a**, Time-lapse acquisition of 100 nm fluorescent beads moved on a motion stage at approximately 50 mm/s to emulate continuous flow. Scale bar, 5 μm. **b**, WF, flow-axis SIM, and SR-IFC images of the same beads. Scale bars, 5 μm. **c**, Point-spread functions obtained by averaging single beads extracted from the images in b. Scale bars, 500 nm. **d**, Population-averaged intensity profiles along the flow and transverse axes, giving SR-IFC full width at half maximum (FWHM) values of 155 nm and 149 nm, respectively, or approximately 118 nm and 110 nm after correction for the 100 nm bead diameter. **e**, Time-lapse acquisition of flowing Jurkat cells at 25 mm/s. Scale bars, 5 μm. **f**, WF, flow-axis SIM and SR-IFC mitochondrial images of the same Jurkat cell after registration. Scale bars, 5 μm. **g**, Magnified views of the region outlined by the dashed white box in f. Scale bars, 500 nm. **h**, Intensity profiles across mitochondrial features, showing peaks resolved by SR-IFC but merged by WF imaging.

These bead measurements were performed on a stage, where inter-frame motion was controlled and almost identical for every bead. Continuous flow removes that control: each cell appears at a different, unknown position in each of the three frames, and the displacement varies from cell to cell. Because SIM reconstruction combines these frames as phase-shifted views of the same object, accurate inter-frame displacement estimation is essential. SR-IFC recovers this displacement directly from the data by using frame cross-correlation to measure each cell’s inter-frame translation and small residual rotation, thereby aligning the three frames to a common reference before reconstruction (Methods “Frame registration and blind-SIM reconstruction”; Supplementary Fig. 3 and Supplementary Note 1). We validated this scheme using Jurkat cells stained with MitoTracker Green FM and flowed at 25 mm/s. Three consecutive frames were captured for each moving cell (Fig. 3e), cross-correlation resolved each cell’s inter-frame displacement, and the registered images were reconstructed without artifacts (Fig. 3f). In registered cells, SR-IFC resolved fine mitochondrial fragments and tubular structures that were blurred or merged in the WF and flow-axis SIM images, both across the whole cell and within a zoomed region of interest (Fig. 3g). Line profiles across these features showed peaks that were separated by SR-IFC but unresolved by WF imaging (Fig. 3h). This registration and reconstruction scheme therefore enables organelle-scale imaging under the uncontrolled motion of continuous flow.

Finally, we characterized throughput, which is determined by the field-of-view length, camera frame rate, flow velocity, and the requirement that each cell be sampled across three consecutive phase frames. For the 2048 × 320 pixel readout region used here, the camera operated at approximately 720 frames per second. Because the three phase frames are stepped sequentially, acquisition can be pipelined such that three cells at different positions within the field of view are captured at different points in their three-frame sequences at any given moment. This allows one cell to be completed per frame, giving a theoretical maximum event rate of approximately 720 cells per second. Reaching this ceiling requires a flow velocity that keeps each cell within the field of view for three frames and a sample concentration sufficient to maintain, on average, three cells in the field of view simultaneously. Because the frame rate scales inversely with readout height, throughput is currently camera-limited and trades off against field-of-view size: larger cells requiring taller readout regions are imaged at correspondingly lower rates. Even so, sustaining hundreds of super-resolved events per second places SR-IFC firmly in the cytometric regime, far beyond the throughput of scanning-or localization-based super-resolution microscopy.

### Application to microbiology: population-scale profiling of lipid droplets in *Euglena gracilis*

Having established the resolution and imaging performance of SR-IFC, we next evaluated its ability to quantify organelle morphology across heterogeneous cell populations. We focused on two organelles that present distinct imaging challenges: lipid droplets, which are small, discrete structures whose size, number, and spatial distribution must be resolved, and mitochondria, which form interconnected networks whose morphology must be classified (see the next section). Lipid droplets are dynamic organelles that store neutral lipids and coordinate energy metabolism, and their number, size, and spatial arrangement reflect cellular metabolic state^47^. In microalgae, they also function as carbon-storage organelles whose biogenesis responds sensitively to nutrient stress, making them valuable readouts for metabolic engineering^48,49^. Droplet size, number, and spacing are, however, difficult to measure accurately when the droplets are smaller than, or closer together than, the diffraction limit. We therefore profiled lipid droplets in nitrogen-starved *Euglena gracilis*, in which all three properties changes as lipid biogenesis proceeds. Cells grown under nitrogen-replete conditions were transferred to nitrogen-free medium, sampled at 0, 4, 8, and 16 h, stained with BODIPY 505/515, and imaged by SR-IFC (Fig. 4a; Methods, “*Euglena gracilis* culture and nitrogen deprivation”). In paired images of the same cells, small droplets that SR-IFC resolved as distinct objects were blurred, merged, or undetectable in the WF images (Fig. 4b, arrows), a difference that was consistent across a large gallery of cells (Supplementary Fig. 4).

**Fig. 4.**
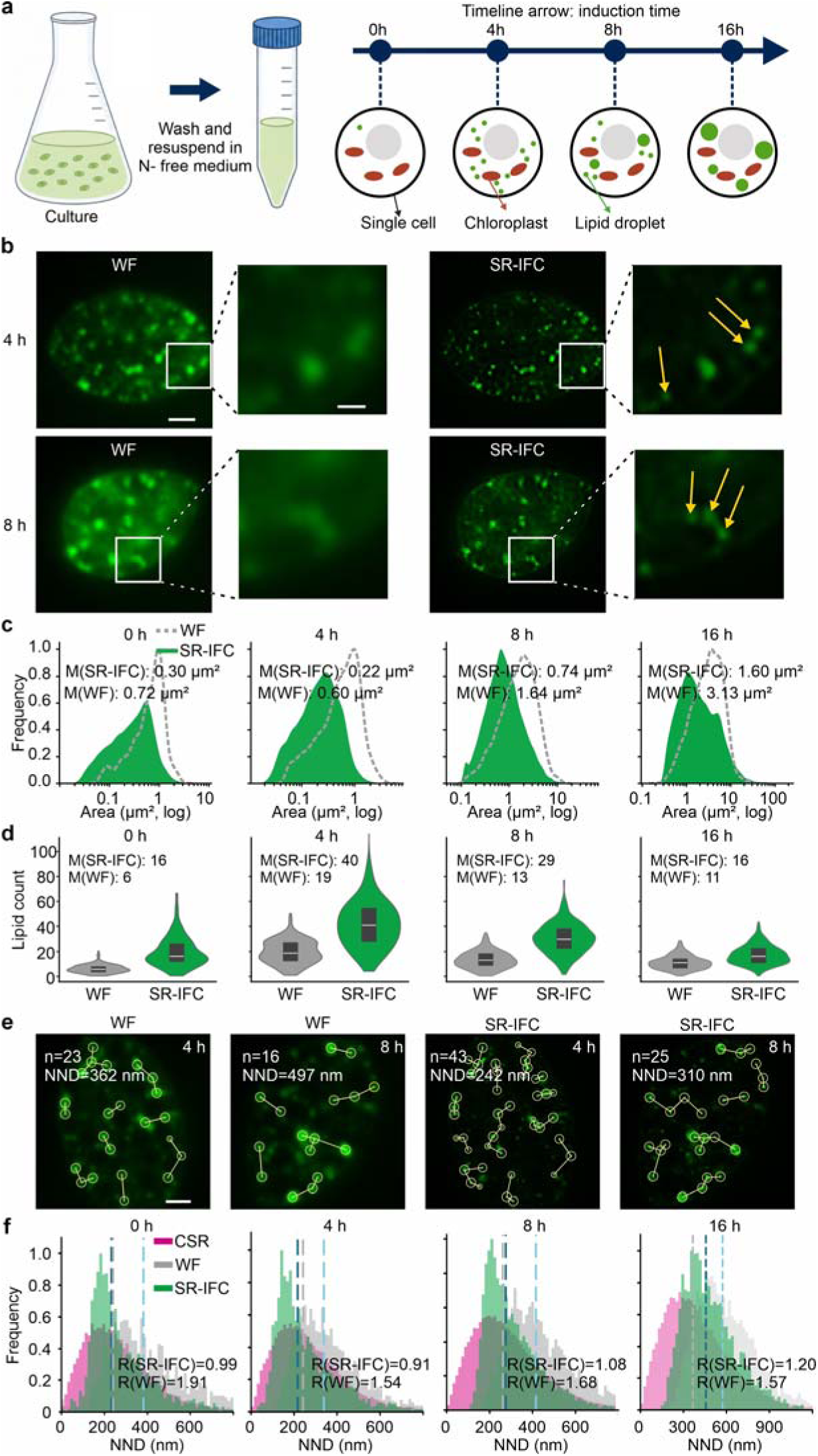
Application to microbiology: population-scale profiling of lipid droplets in *Euglena gracilis*. **a**, Workflow for nitrogen deprivation showing culture, resuspension in nitrogen-free medium, and SR-IFC imaging at 0, 4, 8, and 16 h, with cartoons indicating progressive lipid-droplet accumulation alongside chloroplasts. **b**, Representative cells at 4 and 8 h after nitrogen removal. For each cell, the WF and SR-IFC images are followed immediately by magnified views of the regions outlined in white. Arrows indicate lipid droplets. Scale bars, 5 μm (whole-cell images) and 200 nm (magnified regions of interest). **c**, Individual-droplet area distributions for WF (dashed) and SR-IFC (filled) across the four time points. Median droplet areas are 0.30, 0.22, 0.74, and 1.60 μm² for SR-IFC and 0.72, 0.60, 1.64, and 3.13 μm² for WF. **d**, Per-cell droplet-count distributions shown as paired WF and SR-IFC violin plots. Deep gray boxes denote interquartile ranges and white lines denote medians. Median counts are 16, 40, 29, and 16 droplets per cell for SR-IFC and 6, 19, 13, and 11 for WF. **e**, Representative single-cell nearest-neighbor distance (NND) maps with droplet outlines for WF and SR-IFC at 4 and 8 h, annotated with the corresponding droplet count and median NND for each cell. Scale bars, 5 μm. **f**, NND distributions for SR-IFC (green), WF (gray), and cell-matched complete spatial randomness (CSR) simulations (magenta). For each cell, R is the observed median NND divided by the CSR-expected median NND; R < 1, R ≈ 1, and R > 1 indicate clustering, random placement, and dispersion, respectively.

Because these differences affect what is counted, we segmented every droplet in the WF and SR-IFC images (segmentation pipeline in Supplementary Fig. 5; see Methods, “Image segmentation and morphological analysis”) and quantified per-cell droplet area (Fig. 4c) and droplet count (Fig. 4d) across the four time points. The higher resolution of SR-IFC substantially changed the quantitative picture. WF systematically overestimated droplet area because its diffraction-limited point spread function broadened small droplets and fused neighboring ones. In contrast, SR-IFC returned smaller, better-resolved area distributions at every time point: the median SR-IFC droplet area was 0.30 μm² at 0 h, decreased to 0.22 μm² at 4 h, and then increased to 0.74 μm² at 8 h and 1.60 μm² at 16 h, compared with inflated WF medians of 0.72, 0.60, 1.64 and 3.13 μm², respectively. Droplet counts showed a complementary trend. SR-IFC revealed a biphasic trajectory, with a median of 16 droplets per cell at 0 h, increasing to 40 at 4 h, and then decreasing to 29 at 8 h and 16 at 16 h. WF undercounted droplets at every time point, with medians of 6, 19, 13, and 11, respectively, most severely at 0 and 4 h, when droplets were smallest and most densely packed. Together, the changes in droplet area and the biphasic pattern in droplet count indicate a three-phase lipid-biogenesis program: nucleation of many small droplets peaking at 4 h, droplet growth with concomitant reduction in number at 8 h, and consolidation into fewer, larger droplets by 16 h. This pattern is consistent with the early proliferation and subsequent coalescence reported for nitrogen-starved microalgae^49,50^. Crucially, SR-IFC measures this balance without systematic bias: WF inflation of droplet area and undercounting of droplet number distort absolute quantification of nucleation, growth, and coalescence when the biology is most active.

Sub-diffraction resolution also enabled a readout that is inaccessible to diffraction-limited IFC: the spatial arrangement of droplets within each cell. For every cell, we computed the nearest-neighbor distance (NND) of each droplet in both modalities. In two representative cells, SR-IFC resolved 43 and 25 droplets with median NNDs of 242 and 310 nm, respectively, whereas WF detected only 23 and 16 droplets with inflated median NNDs of 362 and 497 nm (Fig. 4e). This reflects the same WF merging artifact, now expressed as too few droplets separated by artificially large distances. Raw NND, however, depends on droplet density: the more droplets a cell contains, the smaller the spacing, so raw NNDs cannot be compared directly between cells or between WF and SR-IFC, which detect different droplet numbers. To remove this dependence and isolate the spatial pattern, we referenced each cell’s observed median NND to the median NND expected if the same number of droplets were placed at random within the same cell area, a complete spatial randomness (CSR) null model^51^ (CSR construction in Supplementary Fig. 6). This ratio, *R*, defined as the observed median NND divided by the CSR-expected median NND, has a simple interpretation: 1 means the droplets are arranged as if placed at random, 1 means they are more regularly spaced (dispersed, mutually avoiding) than random, and 1 means they are clustered. Because *R* is referenced to each cell’s own random expectation, it makes the WF and SR-IFC arrangements directly comparable despite their different droplet counts. In this way, the two modalities show opposite stories (Fig. 4f; see Methods, “Spatial point-pattern analysis of lipid droplets”). For SR-IFC, the median R varied from 0.99 at 0 h to 0.91, 1.08 and 1.20 at 4, 8 and 16 h, essentially random early, then progressively dispersed, capturing a real reorganization in which droplets coalesce into fewer, larger, mutually excluding bodies. For WF, *R* was high and essentially time-invariant (1.91, 1.54, 1.68 and 1.57) because WF merges unresolved close neighbors into single objects, it preferentially eliminates the shortest inter-droplet distances, inflating the apparent spacing above the random expectation at every time point and producing a spurious, constant “dispersed” signal. Diffraction-limited imaging therefore does not merely add noise to this measurement; it replaces a real temporal reorganization with a persistent artifact.

Taken together, the droplet positions and spacings resolved by SR-IFC indicate that lipid droplets undergo coordinated spatial reorganization during maturation, in addition to changes in number and size. Because lipid-droplet biogenesis and growth are known to nucleate at endoplasmic reticulum (ER) contact sites^52,53^, adding an ER marker could place this reorganization directly within its membrane context. More broadly, these results show that diffraction-limited imaging biases not only droplet size and count but also their spatial relationships, thereby distorting inferences about the timing of nucleation, growth, and lipid trafficking. SR-IFC, in contrast, recovers all three features at population scale. These findings position SR-IFC as a route to image-defined, population-scale readouts of lipid-droplet biology, with applications in metabolic-engineering and biofuel screens in algae, lipid-trafficking disease models, and stress-response phenotyping across single-celled and multicellular systems^19^.

### Application to immunology: phenotyping of mitochondrial morphology in activated CD8^+^ T cells

The second organelle, the mitochondrion, poses a different measurement problem: rather than discrete objects to be counted, it forms a connected, dynamic network whose morphology must be classified, and whose cell-to-cell differences in connectivity become apparent only when individual tubules and their branch points are resolved. Mitochondria undergo extensive remodeling as CD8^+^ T cells activate and differentiate. Naive, memory, and effector states exhibit distinct mitochondrial morphologies, including fused, elongated networks in memory cells and fragmented, punctate mitochondria in effector cells, reflecting the oxidative and glycolytic programs that support these states^54–57^. Mitochondrial morphology may therefore serve as an intrinsic, surface-marker-independent correlate of T-cell state. However, this relationship has been characterized primarily through low-throughput confocal and electron microscopy of individual cells^58,59^, and diffraction-limited imaging flow cytometry cannot resolve the fine architecture on which the distinction rests. We therefore used SR-IFC to read mitochondrial morphology across a large population of cells at sub-diffraction resolution. Naive CD8^+^ T cells were isolated from a healthy donor and subjected to repeated anti-CD3/CD28 stimulation^60^, with samples collected at baseline and after 1, 2, and 3 rounds of stimulation, each lasting 48 h (Fig. 5a; Methods, “Human CD8^+^ T-cell isolation and stimulation”). With mitochondria labeled by MitoTracker Green, SR-IFC resolved structures that appeared blurred or merged in the corresponding WF images (Fig. 5b). The reconstructions also revealed pronounced heterogeneity in mitochondrial architecture, including compact, punctate, elongated, and network-like forms, even within the same stimulation condition (Supplementary Fig. 7).

**Fig. 5.**
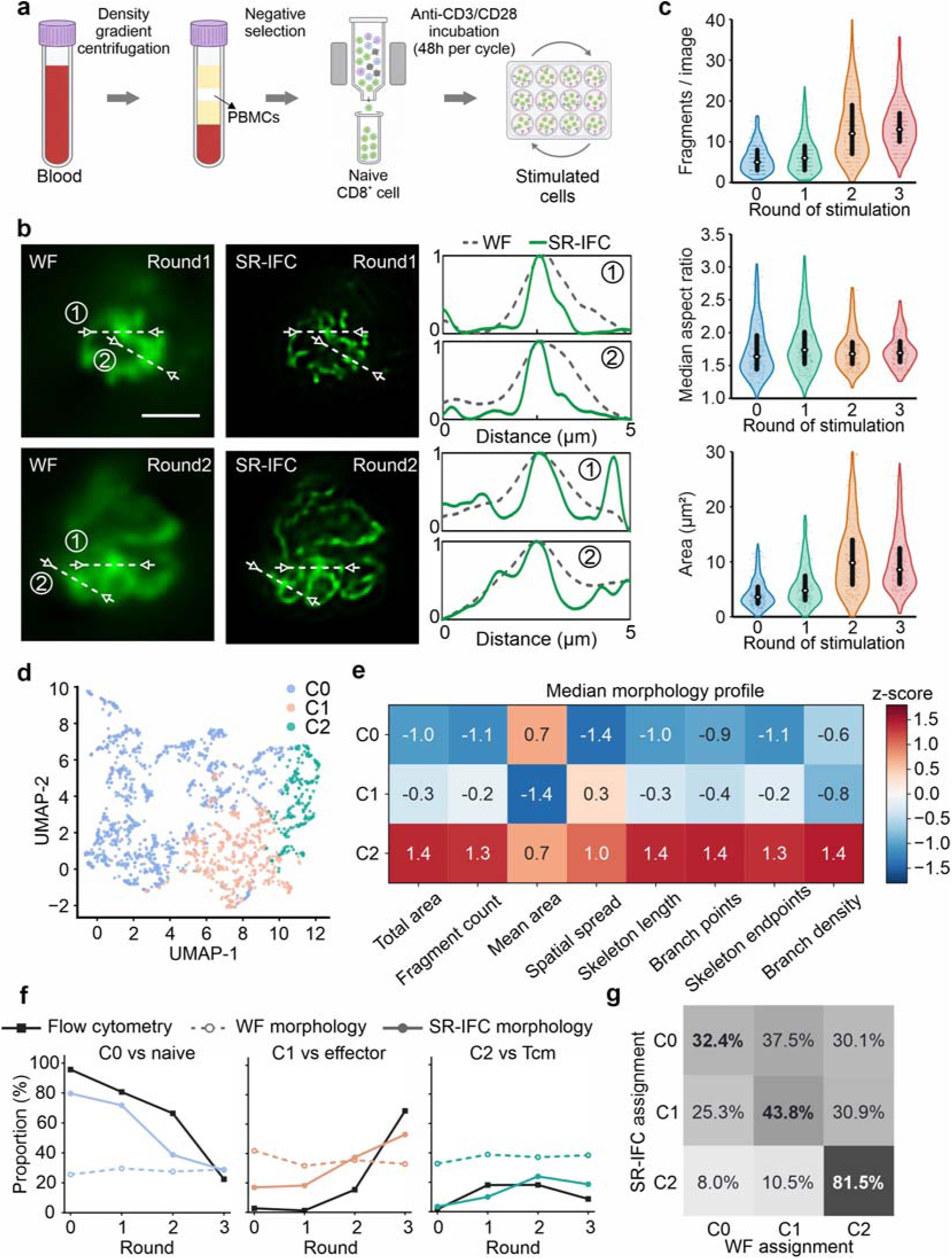
Application to immunology: phenotyping of mitochondrial morphology in activated CD8^+^ T cells. **a**, Workflow showing density-gradient centrifugation, PBMC isolation, negative selection of naive CD8^+^ T cells, and repeated anti-CD3/CD28 stimulation (48 h per round). **b**, WF and SR-IFC mitochondrial images of representative CD8^+^ T cells, with intensity profiles along the two labeled white dashed lines showing peaks resolved by SR-IFC but merged in WF. Scale bar, 5 μm. **c**, Classical morphometrics across 0, 1, 2, and 3 stimulation rounds: fragment count, median fragment aspect ratio, and total mitochondrial area. **d**, UMAP of the self-supervised embeddings colored by k-means cluster (C0 to C2). **e**, Median morphometric profile of each morphotype for eight features: total area, fragment count, mean fragment area, spatial spread, skeleton length, branch points, skeleton endpoints, and branch density. For each feature, the three morphotype medians are standardized across C0 to C2 and shown as z-scores. **f**, Round-resolved fractions of SR-IFC morphotypes (colored solid lines), WF-native morphotypes (colored dashed lines with open markers), and matched flow cytometric phenotypes (black solid lines with square markers): C0 versus naive, C1 versus effector (Tem and TEMRA pooled), and C2 versus Tcm. **g**, Per-cell agreement between the morphotype assignment of the SR-IFC and WF-native encoders, trained independently with the same workflow on matched images. Rows denote SR-IFC morphotypes, columns denote matched WF clusters, and values are row-normalized percentages.

Classical scalar morphometrics captured broad trends during stimulation. Fragment count increased, aspect ratio changed only modestly, and mitochondrial area increased through the second round before reaching a plateau (Fig. 5c). These trends are consistent with the established relationships among CD8^+^ T-cell differentiation, mitochondrial remodeling, and metabolic reprogramming^54,61^. However, because these scalar metrics comprise a limited set of predefined, population-averaged statistics, they capture overall trends but obscure coexisting morphotypes within each stimulation round, potentially averaging out transient minority states. To preserve this information and capture the transient, highly networked intermediate that emerges midway through stimulation, we profiled the SR-IFC images with a self-supervised learning model that requires no manual annotation. A ConvNeXt network^62^, trained with a SimCLR contrastive objective^63^, mapped each single-cell image to a 768-dimensional embedding. The resulting embeddings were clustered using k-means and visualized by uniform manifold approximation and projection (UMAP; Fig. 5d, Supplementary Fig. 8). Across K values from 3 to 6, K = 3 yielded the highest Calinski-Harabasz score, indicating the strongest between-cluster separation relative to within-cluster dispersion. It was also the most reproducible under subsampling and the most readily recovered from independent morphometric features. Although the silhouette score marginally favored a higher K, this was not accompanied by improved stability or recoverability (Supplementary Fig. 9; Methods, “AI phenotyping pipeline”). We therefore selected K = 3 for subsequent analysis, as clustering at K = 4 to 6 only subdivided regions that the three clusters already covered (Supplementary Fig. 10). Morphometric characterization of the same cells identified the three clusters as a sparse morphotype (C0, few mitochondria and low mitochondrial content), a fragmented morphotype (C1, numerous small, poorly connected mitochondria), and a fused morphotype (C2, an extended, highly branched mitochondrial network) (Fig. 5e; Supplementary Fig. 11). The fragmented and fused morphotypes (Supplementary Fig. 12) were consistent with the punctate and fused mitochondrial structures previously reported in effector and memory CD8^+^ T cells, respectively^54^.

The proportions of these morphotypes changed systematically with repeated stimulation (Fig. 5f, solid lines; Supplementary Fig. 13). The sparse morphotype decreased from 79.7% to 28.5%, whereas the fragmented morphotype increased from 16.9% to 52.9% and became dominant by the third round. The fused morphotype increased transiently, reaching a peak of 24.0% after the second round before declining. Cell density shifted continuously across the embedding space between stimulation rounds (Supplementary Fig. 13), indicating a graded redistribution along the morphology manifold rather than abrupt transitions among discrete states. To compare this image-defined trajectory with canonical T-cell differentiation, we performed matched flow cytometric analysis of CD45RA and CCR7^64,65^ using cells from the same batch (Supplementary Fig. 14), resolving naive (CD45RA^+^CCR7^+^), central-memory (Tcm, CD45RA^−^CCR7^+^), effector-memory (Tem, CD45RA^−^CCR7^−^) and TEMRA (CD45RA^+^CCR7^−^) populations as successive stages of CD8^+^ T-cell differentiation^66,67^. We pooled Tem and TEMRA populations into a single effector fraction because both represent effector-differentiated states characterized by similarly fragmented mitochondrial morphologies that were not expected to be distinguishable by morphology alone^54,66,67^. The marker-defined population followed the expected trajectory: the naive fraction declined from 95.7% to 22.4%; the Tcm fraction rose to 18.1% after the first round, remained at 18.2% after the second, and declined to 8.6% after the third; and the pooled effector fraction increased to 69.0% by the third round. Each morphotype defined by SR-IFC covaried with its corresponding marker-defined population (Fig. 5f). We quantified this agreement with Pearson’s correlation coefficient (r), the concordance correlation coefficient (CCC), and the root-mean-square error (RMSE), as shown in Supplementary Fig. 15d. Unlike r, CCC and RMSE also test absolute agreement rather than trend alone. The sparse morphotype tracked the naive fraction (r = 0.90, CCC = 0.81), and the fragmented morphotype tracked the pooled effector fraction (r = 0.93, CCC = 0.73), with RMSEs of 16.9 and 17.6 percentage points, respectively. The fused morphotype also rose and fell with the Tcm fraction, but it rose more slowly and peaked slightly higher, at 24.0% after the second round, so its correlation and concordance were weaker (r = 0.60, CCC = 0.57); its RMSE (7.1 percentage points) was the smallest of the three pairs only because both fractions remained small. These results, obtained in a single donor, show that the sparse and fragmented morphotypes track the naive and effector fractions in both trend and magnitude, so that mitochondrial morphology reads the main axis of CD8□ T-cell differentiation without surface markers and complements marker-based phenotyping. The relatively weak match between the fused morphotype and the central-memory fraction indicates that the transient networked state is not simply equivalent to the Tcm gate, and its relationship to central memory requires further validation.

We next examined whether reading this trajectory requires super-resolution. To provide a fair comparison, we independently trained the same self-supervised model on matched WF images of the same cells, using identical architecture, augmentations, and clustering procedure. This allowed each imaging modality to be analyzed using its own trained encoder within an otherwise matched workflow (Supplementary Fig. 15). The resulting WF-native morphotypes were less clearly separated (Supplementary Fig. 15a,b), and their proportions remained nearly unchanged across the four stimulation rounds (Fig. 5f, dashed lines; Supplementary Fig. 15c), failing to reproduce the decline in the sparse morphotype, the expansion of the fragmented morphotype, or the transient peak of the fused morphotype observed by SR-IFC. Consistent with this result, the WF-derived fractions showed poor correspondence with the matched flow cytometric fractions. Pearson’s r values were −0.52, −0.38, and 0.76, and CCC values were −0.02, −0.09, and 0.05, for the naive, effector, and Tcm comparisons, respectively (Supplementary Fig. 15d). The r value was negative for the naive and effector comparisons, and although the Tcm comparison showed a positive trend, its CCC was near zero because the WF-derived fused fraction (32.6% to 38.8%) exceeded the Tcm fraction by 19 to 31 percentage points in every round. A per-cell comparison further revealed the basis of this discrepancy (Fig. 5g). The fused morphotype was recovered with 81.5% agreement, whereas agreement for the sparse and fragmented morphotypes was only 32.4% and 43.8%, respectively. Approximately one-third of the cells classified as sparse or fragmented by SR-IFC were instead assigned to the fused morphotype by WF. This is expected consequence of a diffraction-limited point-spread function, which merges small, closely spaced mitochondria into apparently connected networks. Consequently, WF overestimated the fused morphotype and obscured the differentiation-associated dynamics of the sparse and fragmented morphotypes. Because these limitations persisted when the WF images were analyzed using an independently trained WF-native encoder, they cannot be explained solely by domain mismatch and instead support an intrinsic limitation of diffraction-limited resolution. The connectivity-based distinctions among sparse, fragmented, and fused mitochondria, including the transient fused intermediate, were therefore not reliably resolved by diffraction-limited imaging but became apparent with population-scale super-resolution. By profiling mitochondrial morphology at the single-cell level, SR-IFC resolved a coordinated remodeling trajectory together with the heterogeneity and continuity obscured by population averaging and two-marker gating. These findings position mitochondrial morphology with SR-IFC analysis as a surface-marker-independent axis for single-cell immunophenotyping, with potential applications in immunometabolism and in monitoring T-cell differentiation during cell manufacturing and adoptive cell therapy.

## DISCUSSION

We developed SR-IFC, a high-throughput platform that reconstructs isotropic in-plane super-resolution images of continuously flowing cells from three phase-shifted frames per cell. The system achieves an in-plane resolution of approximately 118 nm, representing an approximately twofold improvement over the diffraction limit, at throughputs of up to approximately 720 cells per second. SR-IFC overcomes the incompatibility between structured illumination and continuous flow through the coordinated operation of several modules. Stroboscopic excitation preserves fringe contrast during cell motion, three-dimensional hydrodynamic focusing confines cells within the depth of field, and single-orientation SIM physically captures high-spatial-frequency information along the flow axis. A transverse-axis network then reconstructs the corresponding resolution enhancement along the orthogonal axis, which conventional SIM would require additional grating orientations to obtain. Previous studies have combined image-scanning, structured-illumination or light-sheet microscopy with microfluidic systems to achieve sub-diffraction imaging^28^, but these approaches have generally provided limited or predominantly axial resolution enhancement at sub-cytometric rates. To our knowledge, SR-IFC is the first flow-imaging platform to achieve isotropic in-plane super-resolution at throughputs of hundreds of cells per second.

In two biological applications, SR-IFC converted organelle morphology into population-scale quantitative readouts that diffraction-limited imaging compressed or misrepresented. In nitrogen-starved *Euglena gracilis*, it resolved individual lipid droplets and revealed spatial reorganization during lipid-droplet biogenesis that diffraction-limited imaging replaced with a persistent artifact. In primary human CD8□ T cells, it read activation-associated mitochondrial remodeling as a surface-marker-independent trajectory of differentiation, from sparse mitochondria in naive cells, through a transient networked state that peaked after the second stimulation round, to fragmented mitochondria in effector cells. This trajectory could not be recovered from diffraction-limited images even with an independently trained model, so the finding depends on the structural information that super-resolution adds rather than on the analysis pipeline. The trajectory also agrees with previous evidence that mitochondrial architecture is actively remodeled during T-cell differentiation. Memory CD8□ T cells exhibit elongated, interconnected mitochondrial networks, whereas effector T cells display punctate, fragmented mitochondria^54^. OPA1-dependent fusion is required for efficient memory formation and supports cristae organization and oxidative metabolism^54^, whereas DRP1-dependent fission supports the metabolic reprogramming, clonal expansion, migration and tissue infiltration of effector cells^68^. In addition, mitochondrial positioning sustains Ca²□ signaling at the immunological synapse during activation^69^. The transient fused morphotype and the later fragmented morphotype observed here are consistent with these fusion- and fission-dominated phases. Mitochondrial remodeling also accompanies T-cell exhaustion, the dysfunction state that develops under chronic antigen stimulation: progenitor exhausted T cells, which retain the capacity to respond to immune checkpoint blockade, preserve mitochondrial fitness^70^, whereas terminally exhausted cells show disturbed mitochondrial dynamics and accumulate depolarized mitochondria^71^. Because our cells were stimulated repeatedly, part of the fragmented morphotype may reflect exhaustion-related remodeling. As exhaustion markers were not assessed here, establishing this relationship with SR-IFC is an important direction for future work.

Several limitations remain in the current implementation. Resolution enhancement has been demonstrated in a single-color fluorescence channel and in two dimensions, and physical structured illumination is applied only along the flow axis. The achievable throughput is coupled to the camera region of interest, resulting in a trade-off between imaging area and acquisition speed. In addition, the method requires accurate registration of the three phase-shifted frames, which constrains the maximum usable flow velocity. These limitations define several directions for further development of the method. Multicolor acquisition would enable simultaneous visualization of multiple organelles and their contact sites, whereas axial modulation could extend the isotropic resolution enhancement into three dimensions.

Beyond lipid droplets and mitochondria, the same framework could be applied to any fluorescently labeled organelle, including the endoplasmic reticulum, Golgi, lysosomes, and nucleoli, enabling organelle-resolved studies of secretion, autophagy, and stress responses. Integration with image-activated cell sorting^72,73^ would further allow cells selected by organelle phenotype to be recovered for molecular validation across broader donor populations and cell types. By bridging super-resolution microscopy and population-scale cytometry, SR-IFC provides a scalable framework for image-based profiling of organelle heterogeneity, with potential applications in organelle biology, immunometabolism, metabolic engineering, and single-cell phenotyping.

## METHODS

### Ethics statement

All experiments using human peripheral blood were conducted with approval by the Institutional Ethics Committee of the University of Tokyo School of Medicine (approval no. 11049), in compliance with the relevant guidelines and regulations. Written informed consent was obtained from the healthy participant. No animal experiments were performed in this study.

### Optical system and synchronization

SR-IFC was built around an inverted fluorescence microscope equipped with a 100× oil-immersion objective (UPLSAPO100XO, Olympus, NA 1.4) as shown in Fig. 1a. Excitation was provided by a 488 nm continuous-wave diode-pumped laser (Cobolt 06-MLD, Huebner Photonics) coupled into a structured-illumination module containing a 600 grooves/mm visible transmission diffraction grating (GT25-06V, Thorlabs). The grating was oriented perpendicular to the sample-flow direction and mounted on a piezoelectric transducer (PZT; PAS009, Thorlabs) driven by an open-loop piezo controller (MDT694B, Thorlabs). In each acquisition cycle, the PZT stepped the grating through three phase positions separated by one third of the grating period. An order-selection mask blocked the zeroth diffraction order and transmitted the ±1 diffraction orders, which interfered to generate the structured-illumination pattern. Fluorescence was collected through the same objective, separated by a dichroic mirror (Di02-R488-25x36, Semrock), filtered by a bandpass emission filter (FF03-525/50-25, Semrock), and recorded with an ORCA-Flash4.0 sCMOS camera (Hamamatsu, Camera Link interface) operated with experiment-specific regions of interest (ROIs): 2048 × 320 pixels at 720 frames per second for the fluorescent-bead, Jurkat-cell, and CD8^+^ T-cell experiments, and 2048 × 600 pixels at approximately 300 frames per second for the *Euglena* lipid-droplet experiments. The effective pixel size at the sample plane was 65 nm. Camera exposure, PZT actuation, and laser gating were synchronized by a signal generator. For each frame, the PZT step was completed before the laser was gated on for a 300 ns illumination pulse. The laser power at the sample plane, measured after the objective, was approximately 200 mW.

### Microfluidic device

Cells were flowed through a microfluidic chip fabricated by standard soft lithography. The device consisted of a PDMS channel layer replicated from an SU-8 master and irreversibly bonded to a thin coverglass substrate after oxygen-plasma activation. The main channel was 50 μm wide and 35 μm high, with an inlet length of 2 cm and a main-channel length exceeding 4 cm. Three-dimensional hydrodynamic focusing was achieved by introducing sample, side sheath, and top sheath streams at a flow-rate ratio of 1:4:28 using a 20% (w/w) aqueous glycerol solution (viscosity 2.81 cP) as the sheath fluid. At this flow-rate ratio, the sample stream was confined to a region approximately 8 μm thick along the optical axis, centered on the imaging plane. Because this thickness is comparable to the cell diameter, cells occupied a single axial layer, which increased the fraction of cells intersecting the focal plane during acquisition.

### Frame registration and blind-SIM reconstruction

Each cell transiting the structured-illumination field was recorded in three sequential frames, each at a different nominal phase offset of the sinusoidal excitation pattern. Before reconstruction, the frames were aligned to a common reference by cross-correlation-based rigid registration. Inter-frame translation was estimated from the peak of the Cartesian Fourier cross-correlation map and refined to subpixel precision by local peak fitting; the registration procedure also corrected small residual in-plane rotations (Supplementary Fig. 3 and Supplementary Note 1). The WF image of each cell was obtained by averaging the three registered frames, which cancels the sinusoidal illumination pattern and yields a diffraction-limited image acquired with the same photon budget as the reconstructions. To recover the flow-axis super-resolved image (flow-axis SIM) along the physically modulated axis, we applied a filtered blind-SIM algorithm^39^ that simultaneously reconstructs the sample and estimates the illumination patterns without requiring the precise phase offsets a priori. This choice was motivated by the residual phase uncertainty of the PZT-driven grating in the flow-imaging configuration, which can violate the exact phase steps assumed by standard three-phase SIM reconstruction algorithms^38^. The complete derivation of this single-orientation SIM reconstruction is provided in Supplementary Note 2. Fifteen blind-SIM iterations were run per cell to produce the flow-axis SIM image, which extends the spatial-frequency bandwidth along the modulation axis while the orthogonal transverse axis remains at the diffraction limit. The registered WF and flow-axis SIM images of each cell were the two inputs to the reconstruction network described next.

### AI-assisted isotropic reconstruction

To extend super-resolution to the transverse axis without additional grating orientations, we developed a dual-input reconstruction network that learns a shared directional resolution-enhancement operator from the WF and flow-axis SIM images of the cells being imaged (Supplementary Fig. 1). Both inputs were retained at their native sampling of 65 nm per pixel. For measurement-derived directional supervision, a 1.5D encoder was trained using three-pixel-high horizontal strips (H = 3) extracted from the paired images (Supplementary Fig. 1a). Each WF strip served as the input, and the corresponding measured flow-axis SIM strip was used as the fixed reference for the flow-axis loss, *L*_x_, thereby preserving the experimentally measured spatial-frequency information along the flow axis. The encoder used one-dimensional convolution kernels of 7 pixels, with an effective receptive field of approximately 25 pixels along each strip, 24 hidden channels, and 12 output feature channels. The complete WF and flow-axis SIM images were then supplied as two input channels to a depth-4 two-dimensional U-Net with 20, 40, 80, and 120 channels and a 240-channel bottleneck (Supplementary Fig. 1b). This branch provided image-level spatial context to regularize the strip-wise directional mapping. A two-dimensional loss, *L*_2D_, further constrained the estimated transverse-axis detail using the surrounding two-dimensional image structure. The directional and two-dimensional branches were then fused to reconstruct the transverse-axis information (Supplementary Fig. 1c). Three-column strips extracted along the transverse axis of the flow-axis SIM image were transposed into the orientation used for training and processed by the shared encoder. The predicted directional residual was added to the transposed flow-axis SIM baseline, and the result was returned to the original image coordinates by inverse transposition within the same forward pass. The reconstructed transverse-axis information was then combined with the measured flow-axis SIM image to generate the final SR-IFC image. Neither nine-frame SIM images nor separately generated super-resolved images were used as supervisory targets; the network was trained exclusively using paired WF and flow-axis SIM measurements obtained from the same cells. The bead, *Euglena*, Jurkat, and CD8^+^ T-cell datasets were processed using the same network architecture, with model weights trained independently for each specimen type. All models were optimized using AdamW for 30 epochs with a batch size of 16.

### Simulation and image-quality metrics

Each phantom was defined as a fluorophore distribution in a single focal plane and imaged using a forward model that applies the structured-illumination pattern for each raw frame, convolved the resulting image with the microscope point-spread function, sampled it on the camera pixel grid, and added an sCMOS noise model incorporating photon shot noise and camera read noise (Supplementary Table 1). Each reconstruction was compared with the corresponding noiseless ground truth projected onto the image plane. The NRMSE was calculated from the pixel-wise intensity differences between each reconstruction and the ground truth, with a value of 0 indicating a perfect match. The MS-SSIM quantified similarities in local luminance, contrast, and structural patterns across multiple spatial scales on a scale from 0 to 1, with 1 indicating identical images. For the Fourier-domain analysis, the NRMSE of the Fourier amplitude was calculated within each radial spatial-frequency bin and averaged over 12 independent realizations for each phantom. The resulting phantom-specific means were then averaged with equal weighting to obtain a single curve for each imaging mode. For the noise-robustness analysis, peak relative shot noise was defined as the ratio of the shot-noise standard deviation to the peak signal intensity.

### Quantification of the resolved feature fraction

To quantify whether each reconstruction recovered predefined resolution targets, we calculated the resolved-feature fraction (RFF) for the bead-pair, ellipsoid, and frequency-pattern phantoms (Supplementary Table 1). Unlike the pixel-wise NRMSE and MS-SSIM metrics, RFF does not compare every reconstructed pixel with the ground-truth image. Instead, the noiseless three-dimensional phantom and its projected ground-truth metadata provided the position, orientation, separation, aspect ratio and spatial period of each test feature. These ground-truth annotations were used to define identical measurement regions in the WF, flow-axis SIM, SR-IFC, and conventional nine-frame SIM images. Unless otherwise stated, a single relative tolerance of ε = 0.10 was applied to all criteria defined in equations (1) to (6) below.

For the bead-pair phantom (P1), each test feature consisted of two fluorescent beads with predefined center positions, separation and orientation. An intensity profile was measured along the line connecting the two ground-truth bead centers. The valley-depth score *D* was calculated as

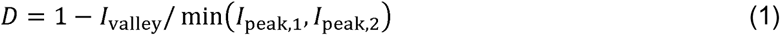

where *I*_peak,1_ and *I*_peak,2_ are the two peak intensities and *I*_valley_ is the minimum intensity between them. A bead pair was considered resolved when two distinct peaks were detected within the two ground-truth-defined bead regions and *D* ≥ ε, that is, when the inter-peak valley lay at least 10% below the smaller peak. A pair was considered unresolved when fewer than two peaks were detected or when *D* < ε.

For the ellipsoid phantom (P2), each elongated test feature had a predefined center, orientation and ground-truth aspect ratio of 1:3, 1:5, or 1:10. The reconstructed major-axis and minor-axis widths (*W*_major_, *W*_minor_) were obtained from the principal axes of the intensity distribution, and the reconstructed aspect ratio *AR*_rec_ was calculated as

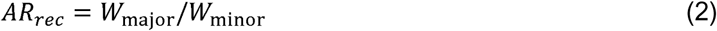

The relative aspect-ratio error *E*_AR_ was calculated as

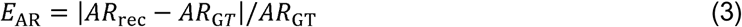

Because an ellipsoid has axial rather than directional orientation, its orientation error Δθ was calculated modulo π:

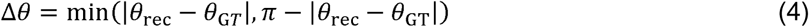

An elongated ellipsoid was considered successfully recovered when *E*_AR_ ≤ ε and Δθ/π ≤ ε. With ε = 0.10, the acceptable reconstructed aspect-ratio ranges were 2.7 to 3.3, 4.5 to 5.5, and 9.0 to 11.0 for ground-truth aspect ratios of 1:3, 1:5, and 1:10, respectively, and the corresponding orientation tolerance was 0.1π rad (18°). Features failing either condition were classified as unresolved. Objects with a ground-truth aspect ratio of 1:1 served as isotropic controls and were considered circularity-preserving when |*AR*_rec_ − 1| ≤ ε; they were excluded from the orientation-recovery calculation because their principal orientation is undefined.

For the frequency-pattern phantom (P3), each test feature consisted of a periodic line-pattern block with a predefined spatial period and orientation. The dominant spatial period and orientation were measured from the two-dimensional Fourier spectrum within the ground-truth-defined block. The relative period error was calculated as

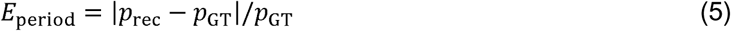

and the axial orientation error was calculated using the same modulo-π definition as for the ellipsoids. Pattern visibility was quantified using the Michelson modulation contrast,

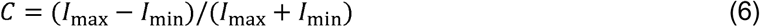

A frequency-pattern block was considered resolved when its relative period error was no greater than ε, its normalized orientation error was no greater than ε and its Michelson contrast was at least ε. Blocks failing any of these conditions were classified as unresolved.

For each phantom, the RFF was calculated as the number of ground-truth-defined features satisfying all corresponding criteria divided by the total number of evaluated features. Features were grouped by separation and orientation for P1, aspect ratio and orientation for P2, and spatial period and orientation for P3. The resolved fractions were first calculated within each group and then averaged with equal weights.

### Image segmentation and morphological analysis

SR-IFC and matched WF images were processed with automated image-analysis pipelines implemented in Python. All measurements were performed at the native pixel size of 65 nm and converted to physical units. For lipid-droplet analysis in *Euglena gracilis*, images were preprocessed by white top-hat background removal, Gaussian smoothing, and percentile-based intensity normalization and enhancement. Lipid-droplet candidates were then segmented by multi-Otsu thresholding, followed by morphological refinement including dilation, closing, hole filling, small-object removal, optional watershed splitting, and shape-based filtering (Supplementary Fig. 5). Extracted metrics included lipid-droplet count, total lipid-droplet area, individual droplet area, integrated intensity, and nearest-neighbor distance.

For CD8^+^ T-cell analysis, individual cell images were first processed with a white top-hat filter using a disk-shaped structuring element to suppress low-frequency background. A coarse foreground mask was generated from non-zero top-hat pixels, followed by binary closing, binary dilation, and removal of small objects. Top-hat intensities within the coarse foreground mask were then linearly normalized to an 8-bit range, while pixels outside the foreground mask were set to zero. Final mitochondrial segmentation was performed within this normalized foreground using an Otsu threshold computed from non-zero foreground pixels. Extracted metrics included fragment count, total organelle area, mean and median fragment area, and mean and median fragment aspect ratio. The same processed single-cell images served as inputs to the self-supervised phenotyping described below.

### *Euglena gracilis* culture and nitrogen deprivation

*Euglena gracilis* NIES-48 was obtained from the Microbial Culture Collection at the National Institute for Environmental Studies and cultured in AF-6 medium in culture flasks (20 mL working volume). Cultures were maintained at 25 °C under a 14:10 hour light/dark cycle at approximately 120 μmol photon m ² s ¹. For nitrogen-deprivation experiments, *E. gracilis* cells were pre-cultured in AF-6 medium, transferred to nitrogen-deficient AF-6 medium (AF-6 lacking the nitrogen source), and sampled at 0, 4, 8, and 16 h after nitrogen removal. For lipid-droplet staining, a stock solution of 1 mM BODIPY 505/515 (Thermo Fisher Scientific) in dimethyl sulfoxide containing 1% ethanol was prepared. *E. gracilis* cells at approximately 10 cells mL□¹ were stained with 10 μM BODIPY 505/515 in deionized water, incubated in the dark for 30 min, washed, resuspended in deionized water, and immediately used for imaging.

### Spatial point-pattern analysis of lipid droplets

Lipid-droplet positions were taken as the centroids of the segmented droplets in each cell. For each droplet, the nearest-neighbor distance (NND) to the closest other droplet was computed, and the per-cell NND distribution was summarized by its median. To test for clustering, the observed NNDs were compared with a complete spatial randomness (CSR) null model^51^, in which the same number of droplets was placed independently and uniformly at random within the cell area; the placement was repeated to estimate the CSR-expected median NND (Supplementary Fig. 6). A per-cell *R* ratio was defined as

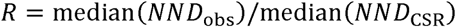

where *R* < 1 indicates clustering, *R ≈* 1 random placement, and *R* >1 dispersion.

### Human CD8^+^ T-cell isolation and stimulation

Human CD8^+^ T cells were isolated from whole blood collected from one healthy male donor. Blood was collected in heparinized vacuum blood collection tubes (Venoject II, Terumo) and processed at room temperature within the recommended hold time. PBMCs were isolated by density-gradient centrifugation over Histopaque-1077 (Sigma-Aldrich, Cat# 10771-500ML) according to the manufacturer’s instructions. The PBMC layer at the interface was collected, washed in PBS, and resuspended for subsequent selection. Naive CD8^+^ T cells were enriched from PBMCs by negative magnetic selection using the MojoSort™ Human CD8 Naive T Cell Isolation Kit (BioLegend, Cat# 480045) with the MojoSort Magnet (BioLegend, Cat# 480020) following the manufacturer’s protocol.

For T-cell activation, 96-well U-bottom plates (Thermo Fisher Scientific, Cat# 163320) were coated with anti-CD3 monoclonal antibody (BD Biosciences, Purified NA/LE Mouse Anti-Human CD3, clone OKT3, Cat# 567107) and anti-CD28 monoclonal antibody (eBioscience/Thermo Fisher, clone CD28.2, Functional Grade, Cat# 16-0289-85) by adding 50 μL of PBS containing 0.5 μg of each antibody (10 μg/mL) per well. Plates were incubated at 20 °C for 30 min to immobilize the antibodies. Wells were aspirated and washed once with PBS before cell seeding. Naive CD8^+^ T cells were seeded at 3.0 × 10 cells per well in 200 μL of RPMI 1640 medium supplemented with 10% human AB serum (Thermo Fisher Scientific, Cat# 12181301C), 2% penicillin-streptomycin, and recombinant human IL-2 (PeproTech, Cat# 200-02-100ug) at a final concentration of 30 IU/mL.

Before stimulation, isolated naive CD8^+^ T cells were divided into four groups designated for analysis after 0, 1, 2, or 3 consecutive rounds of anti-CD3/CD28 stimulation. Each round consisted of a 48 h incubation on plate-bound antibodies. At the end of each round, the group reaching its designated endpoint was harvested, while groups requiring further stimulation were transferred to freshly antibody-coated wells containing fresh complete medium supplemented with IL-2 (30 IU/mL) and maintained under the same culture conditions. At each designated endpoint, including round 0, the harvested cells were divided equally between SR-IFC imaging and matched flow cytometric surface-marker profiling.

For SR-IFC imaging, CD8^+^ T cells were stained with MitoTracker Green FM (Thermo Fisher Scientific, Cat# M7514) at 200 nM final concentration in pre-warmed PBS, incubated for 30 min at 37 °C, washed once with pre-warmed PBS, and resuspended in PBS for imaging. Stained cells were introduced into the SR-IFC microfluidic device under the hydrodynamic focusing and imaging conditions described above.

### Flow cytometric analysis of CD8^+^ T-cell differentiation states

Matched same-batch CD8^+^ T-cell samples were analyzed by flow cytometry to define surface-marker differentiation states for comparison with SR-IFC morphology clusters. The antibody panel included eBioscience Fixable Viability Dye eFluor 780 (Thermo Fisher Scientific, Cat# 65-0865-14), FITC anti-human CD8a antibody (BioLegend, Cat# 301006), APC anti-human CD45RA antibody (BioLegend, Cat# 304111), and PE anti-human CD197 (CCR7) antibody (BioLegend, Cat# 353203). Cells were first stained with the Fixable Viability Dye eFluor 780 at a 1:1000 dilution in PBS for 30 min at room temperature in the dark, then washed once with PBS, and stained with the surface-antibody cocktail in PBS (5 μL of each antibody per test of 0.5 × 10□ cells in 100 μL) for 20 min at room temperature in the dark following the manufacturer’s recommendations, with single-stained compensation controls prepared for each fluorophore. Stained cells were acquired on a CytoFLEX LX flow cytometer (Beckman Coulter) and analyzed in FlowJo. Cells were gated using forward- and side-scatter area parameters, with the scatter gate adjusted to account for stimulation-induced changes in T-cell scatter profiles. Viable CD8^+^ cells were then classified as naive (CD45RA^+^CCR7^+^), Tcm (CD45RA^−^CCR7^+^), Tem (CD45RA^−^CCR7^−^), and TEMRA (CD45RA^+^CCR7^−^) (Supplementary Fig. 14). CD45RA and CCR7 gating thresholds were determined with reference to their expression in naive T cells. Measurements were performed at the same stimulation rounds used for SR-IFC imaging.

### AI phenotyping pipeline

Self-supervised phenotyping of mitochondrial morphology was performed on SR-IFC images of CD8^+^ T cells. Single-cell images processed as described in “Image segmentation and morphological analysis” were used as inputs to a ConvNeXt-Tiny^62^ encoder initialized with ImageNet-pretrained weights (ConvNeXt_Tiny_Weights.DEFAULT, torchvision) and trained with a SimCLR contrastive objective^63^. For grayscale input, the RGB stem weights were averaged across the three input channels. The 768-dimensional encoder output was passed through a projection head (768→256→ReLU→128) for the contrastive loss. The SimCLR temperature was 0.5.

Data augmentations included random resized cropping, horizontal and vertical flipping, rotation, affine translation and scaling, and Gaussian blur. The encoder was optimized with AdamW (learning rate, 5 × 10^−4^; weight decay, 1 × 10^−4^) for 50 epochs using a batch size of 64. The learning-rate schedule comprised 5 epochs of linear warmup followed by cosine decay. Unsupervised clustering was performed on the 768-dimensional embeddings using k-means for *K* = 3 to 6 (Supplementary Figs. 8 and 9). UMAP visualization used n_neighbors = 15, min_dist = 0.1, and cosine distance. For the WF comparison, matched WF images were processed with an independently trained WF-native encoder using the same architecture, augmentations, and clustering procedure (Supplementary Fig. 15).

## Data availability

All relevant data supporting the findings of this study are available within the paper and its Supplementary Information.

## Acknowledgements

We thank Kelvin C. M. Lee and Yoav Shechtman for valuable advice on the microfluidic chip and imaging operation. We extend our gratitude to Serendipity Lab for facilitating collaboration opportunities. No generative AI tool was used to produce, analyze, or interpret experimental data.

## Funding

This work was supported by the Japan Society for the Promotion of Science (JSPS) KAKENHI (JP24K17800, JP26K02037); Japan Science and Technology Agency (JST) ASPIRE (JPMJAP2316); The Mazda Foundation; and the Cell Aging Control Endowed Chair at The University of Tokyo. This work was also supported by Japan Society for the Promotion of Science (JSPS) DC2 (JP26KJ0725). T.N.S. received funding from the Deutsche Forschungsgemeinschaft (DFG, German Research Foundation) – Project-ID 533767322 – EXC 3113/1, Cluster for Nucleic Acid Sciences and Technologies – NUCLEATE and through a JSPS Invitational Fellowship. Yi.L. additionally acknowledges financial support from the FoPM-WINGS program of The University of Tokyo.

## Author contributions

Yi.L., T.D. and K.G. conceived the project. Yi.L. and T.D. designed the SR-IFC system, and Yi.L. and H.W. constructed the system and validated its performance. Yi.L., Yu.L., Y.Z., and M.N. prepared the *Euglena gracilis* and CD8^+^ T-cell samples. Yi.L. conducted the SR-IFC experiments and analyzed the resulting images. Yi.L. and T.D. interpreted the experimental data. Yi.L., T.D., and K.G. wrote the paper. T.I. and Y.T. provided the protocol for preparing primary human CD8^+^ T cells and advised on the corresponding experiments. M.N., T.I., Y.T., and T.N.S. provided valuable comments that improved the quality of the work. All authors participated in reviewing the paper.

## Competing interests

K.G. is a shareholder of CYBO, LucasLand, FlyWorks, and NanoTitan. T.D. is a shareholder of NanoTitan. The remaining authors declare no competing interests.

## SUPPLEMENTARY INFORMATION

This document contains 18 supplementary items referenced from the main manuscript, comprising 15 figures (Supplementary Fig. 1 to Supplementary Fig. 15), Supplementary Table 1, and Supplementary Notes 1 and 2.

---

**Supplementary Fig. 1.**
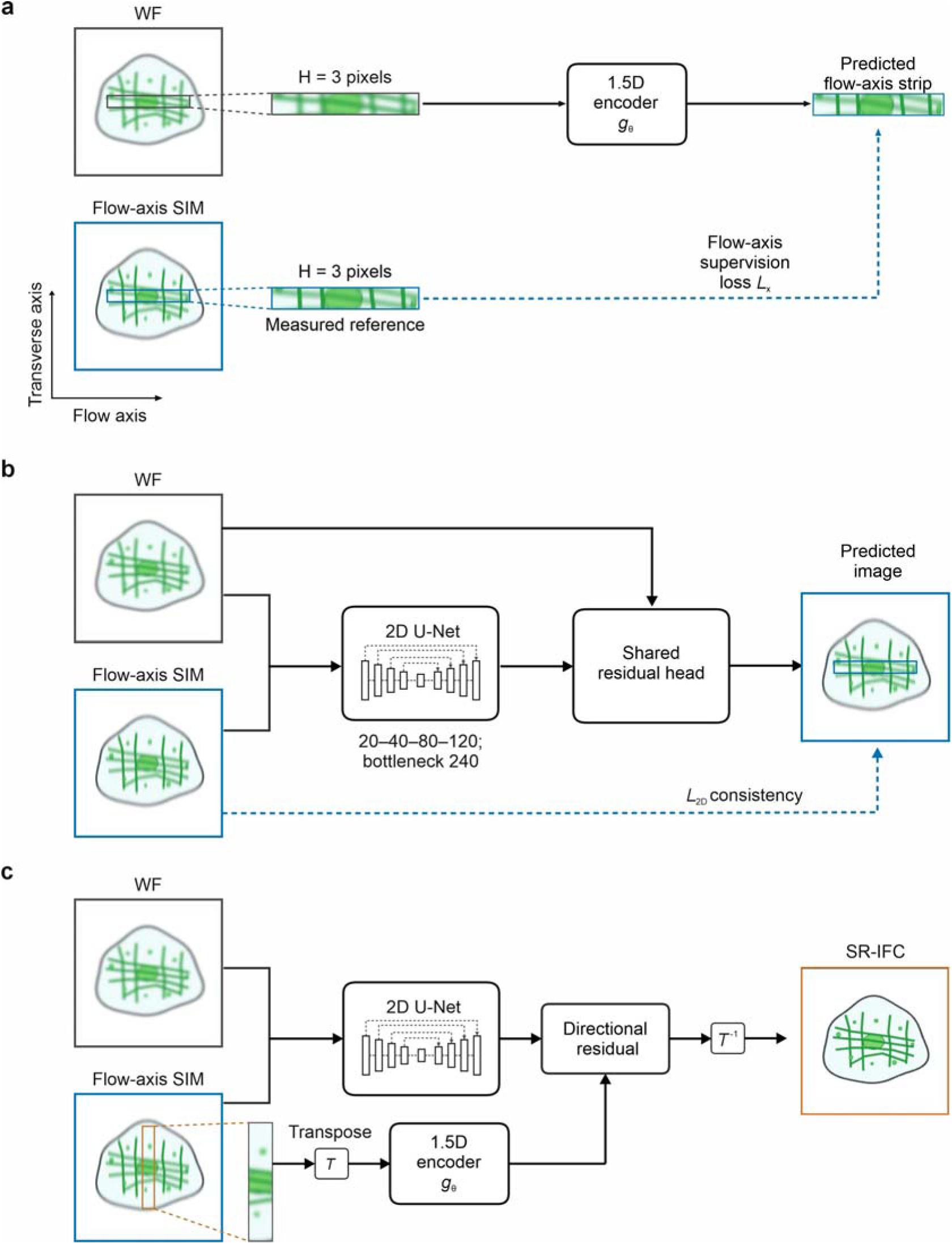
Dual-input U-Net^1^ reconstruction for isotropic in-plane SR-IFC. **a**, A 1.5D encoder *g*_θ_ learns the measured flow-axis resolution enhancement from paired three-pixel-high horizontal strips (H = 3) extracted from registered WF and flow-axis SIM images. The WF strip serves as the input, whereas the corresponding measured flow-axis SIM strip provides the fixed reference for the flow-axis loss *L*_x_. **b**, A two-dimensional U-Net with four downsampling levels, channel widths of 20, 40, 80, and 120, and a 240-channel bottleneck integrates spatial context from the paired WF and flow-axis SIM images. The network predicts spatially varying coefficients for a shared residual head, which combines fixed multiscale filter responses to form a constrained correction to the WF image. The resulting prediction is compared with the paired flow-axis SIM reference through the image-level structural-consistency loss, *L*_2D_, which constrains the estimated transverse-axis detail using the surrounding two-dimensional image structure. **c**, The flow-axis SIM image serves as the restoration input, while the paired WF image provides auxiliary low-frequency context. Transposition, T, reorients the transverse axis to match the direction used during directional learning. The same encoder gθ, two-dimensional U-Net, and residual reconstruction rule are then applied with the restoration input and spatial context reoriented consistently. The predicted residual is added to the transposed flow-axis SIM baseline, and inverse transposition, T^−1^, returns the reconstruction to the original image coordinates to generate the SR-IFC image. The original image dimensions and pixel spacing are retained. Solid black arrows indicate information flow, whereas blue dashed arrows indicate training supervision. Neither conventional nine-frame SIM images nor separately generated super-resolved images are used as supervisory targets.

**Supplementary Fig. 2.**
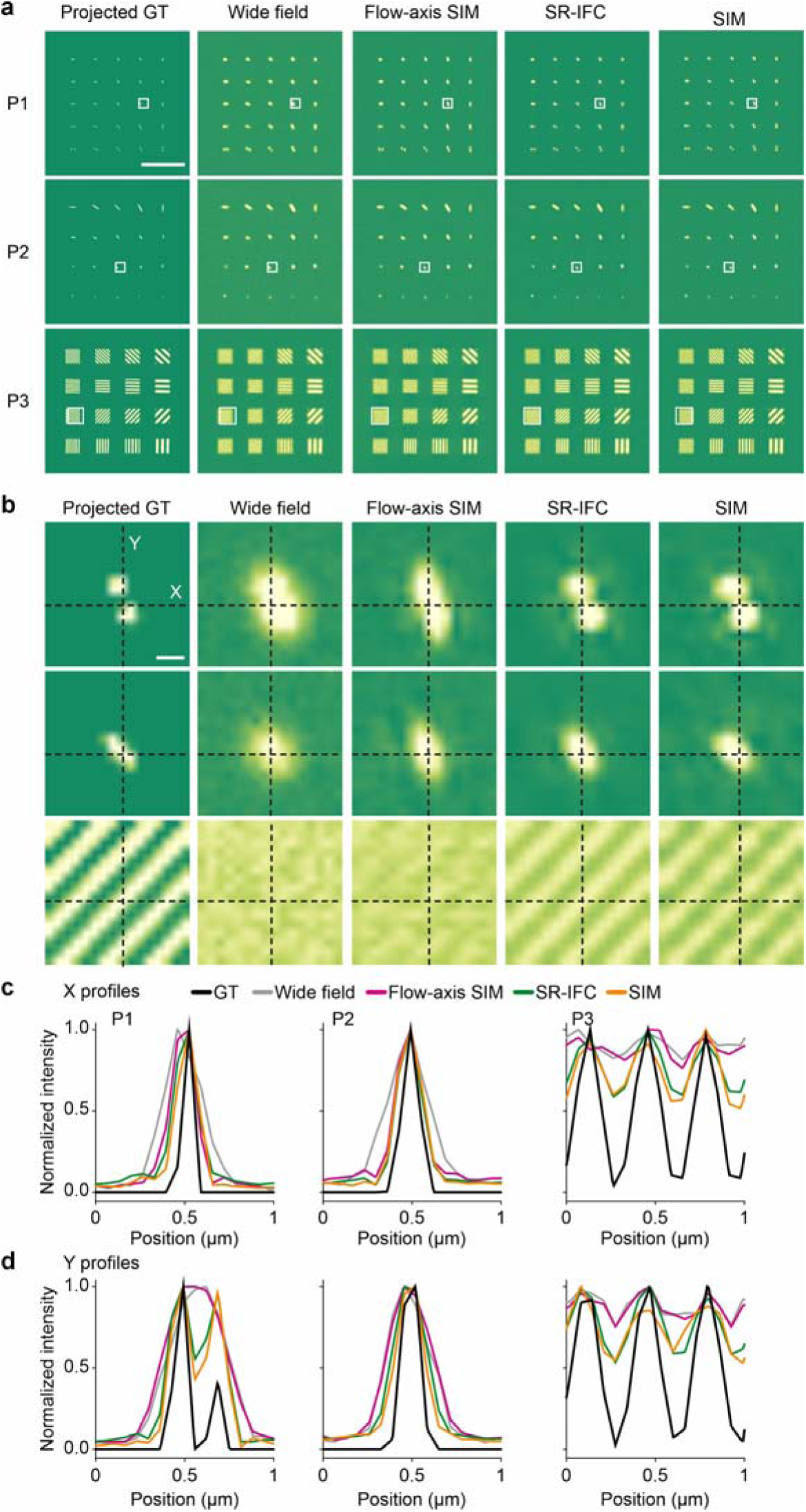
Per-scenario reconstruction comparison with zoomed regions and line profiles. **a**, Bead-pair (P1), ellipsoid (P2), and frequency-target (P3) phantoms over the full field of view, shown as the projected ground truth and as reconstructed by WF imaging, flow-axis SIM, SR-IFC, and nine-frame SIM. Boxed regions are enlarged in **b**. Scale bar, 5 μm. **b**, Magnified regions of interest for each phantom under the four imaging modes, with x and y crosshairs marking the profile locations. Scale bar, 200 nm. **c**, Normalized intensity profiles along x (the flow axis), for each phantom and imaging mode. **d**, Corresponding normalized intensity profiles along y (the transverse axis).

**Supplementary Fig. 3.**
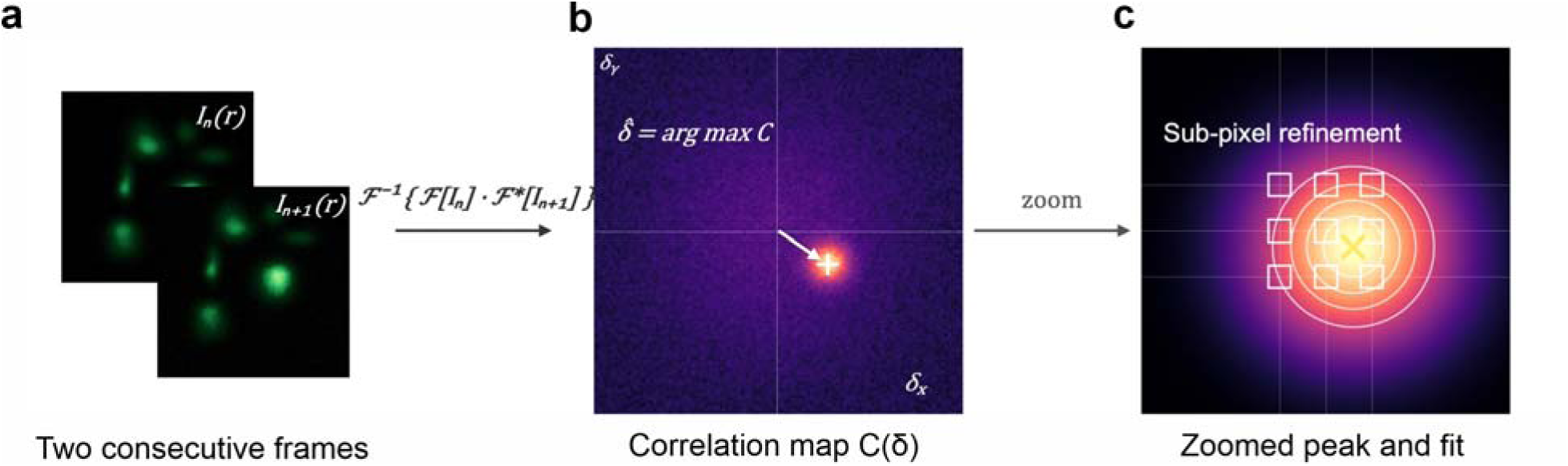
Cross-correlation registration of consecutive flowing-cell frames. Three phase-shifted frames of each cell are acquired stroboscopically at 720 frames per second; within the short inter-frame interval the cell moves to a different position in the camera ROI and may undergo a small angular drift, both of which are corrected before SIM reconstruction by the two-stage (translation, then rotation) procedure described in Supplementary Note 1. **a**, Two consecutive phase frames, *I□*(r) and *I□□□*(r). **b**, Cartesian Fourier cross-correlation map C(δ); its global maximum gives the integer-pixel displacement estimate. **c**, Enlarged correlation peak and local peak fit used for subpixel refinement.

**Supplementary Fig. 4.**
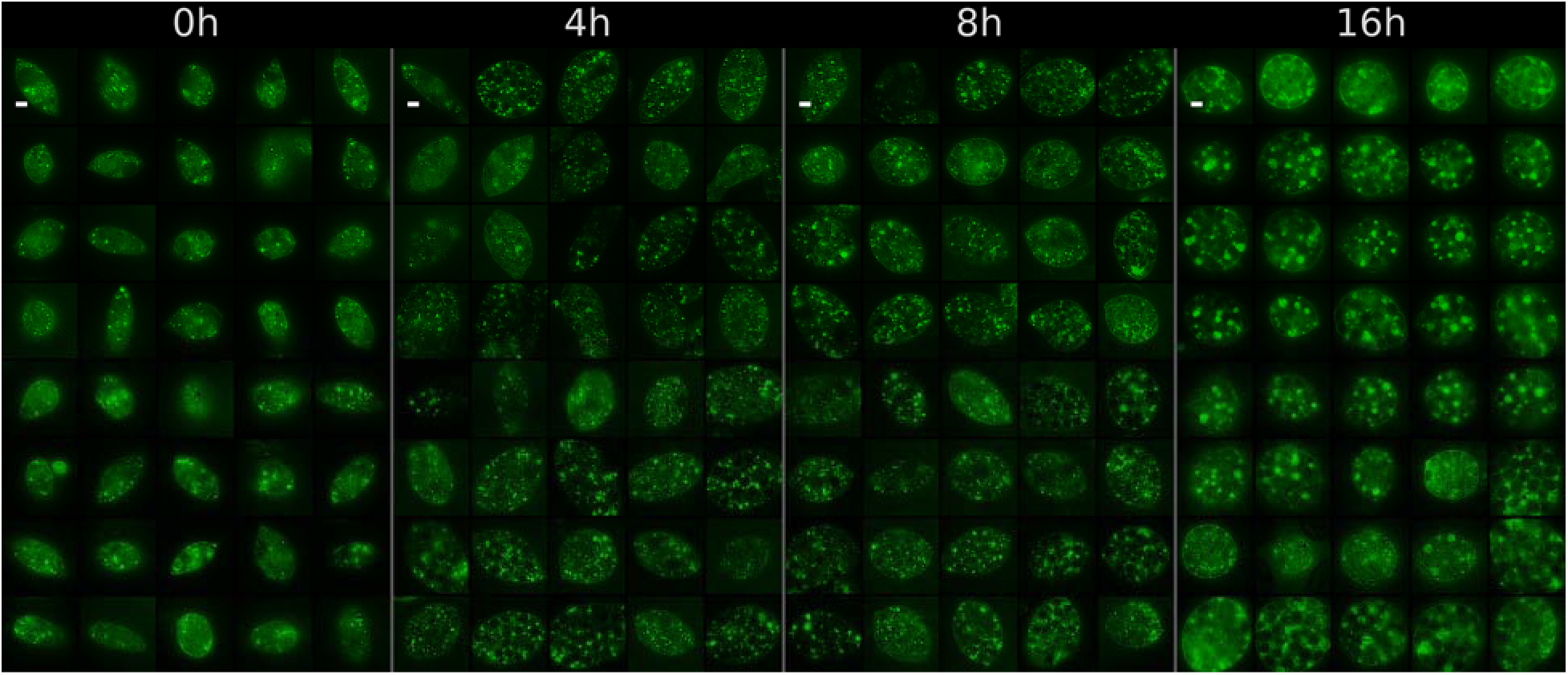
SR-IFC cell gallery for *Euglena gracilis*. Four 5 × 8 montages of randomly sampled SR-IFC single-cell images, one per nitrogen-deprivation time point (0, 4, 8, and 16 h after nitrogen removal), showing the cell-level heterogeneity of intracellular lipid droplets that the population statistics in Fig. 4 summarize. Within a single time point, cells differ markedly in droplet count, droplet size, and spatial distribution. Scale bar, 5 μm.

**Supplementary Fig. 5.**
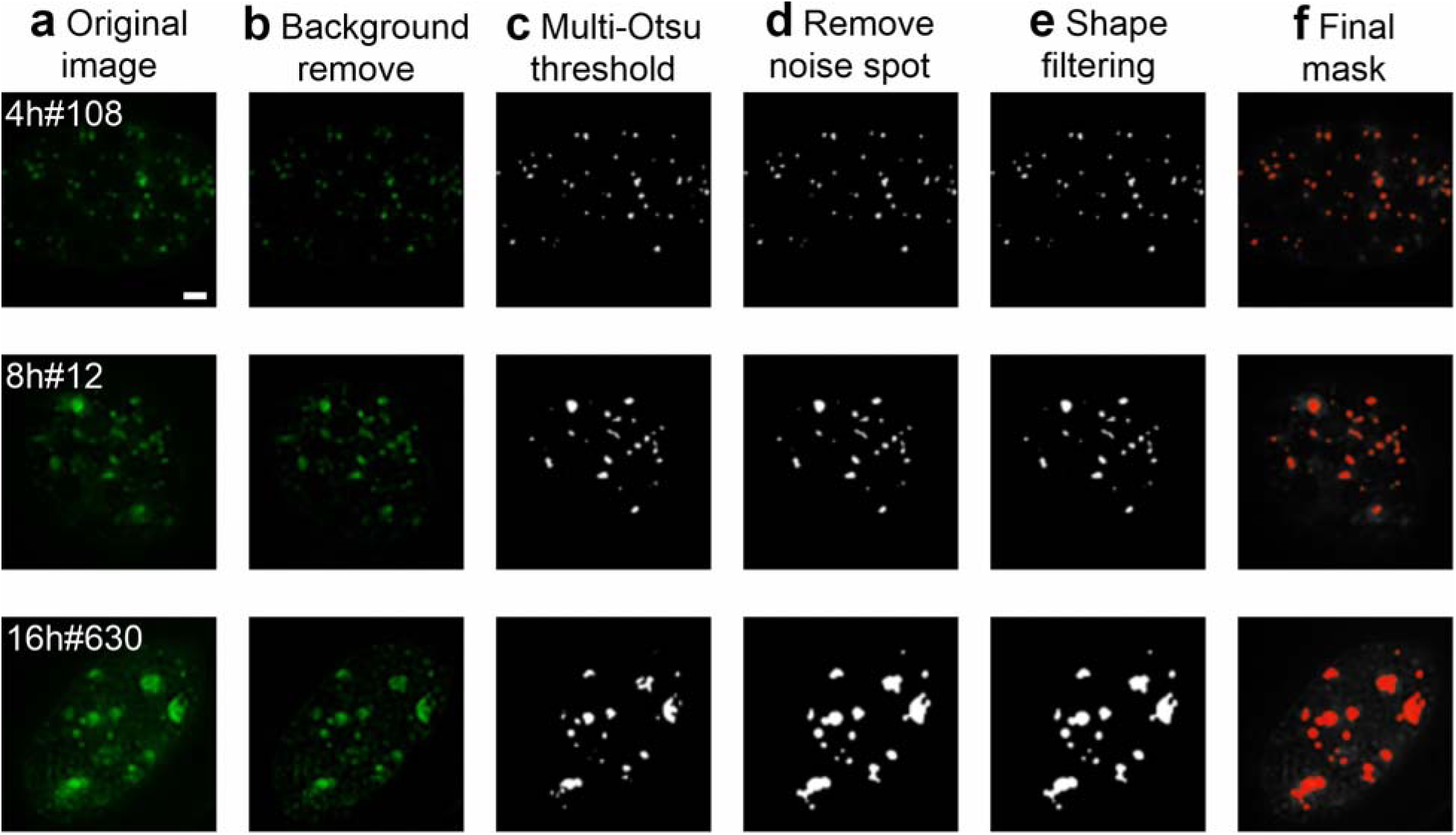
Lipid-droplet segmentation pipeline. Rows show example SR-IFC images acquired at 4, 8, and 16 h after nitrogen removal. The panels show selected stages of the pipeline described in Methods, “Image segmentation and morphological analysis”, and not every operation. **a**, Original image. **b**, Preprocessed image after white top-hat background removal, Gaussian smoothing, and percentile-based intensity normalization and enhancement. **c**, Lipid-droplet candidates after multi-Otsu thresholding. **d**, Candidates after small-object removal; the intervening morphological refinement steps of dilation, closing, hole filling, and optional watershed splitting are applied between c and d and are not shown as separate panels. **e**, Candidates after shape-based filtering. **f**,Final binary mask overlaid in red on the original image. Scale bar, 5 μm.

**Supplementary Fig. 6.**
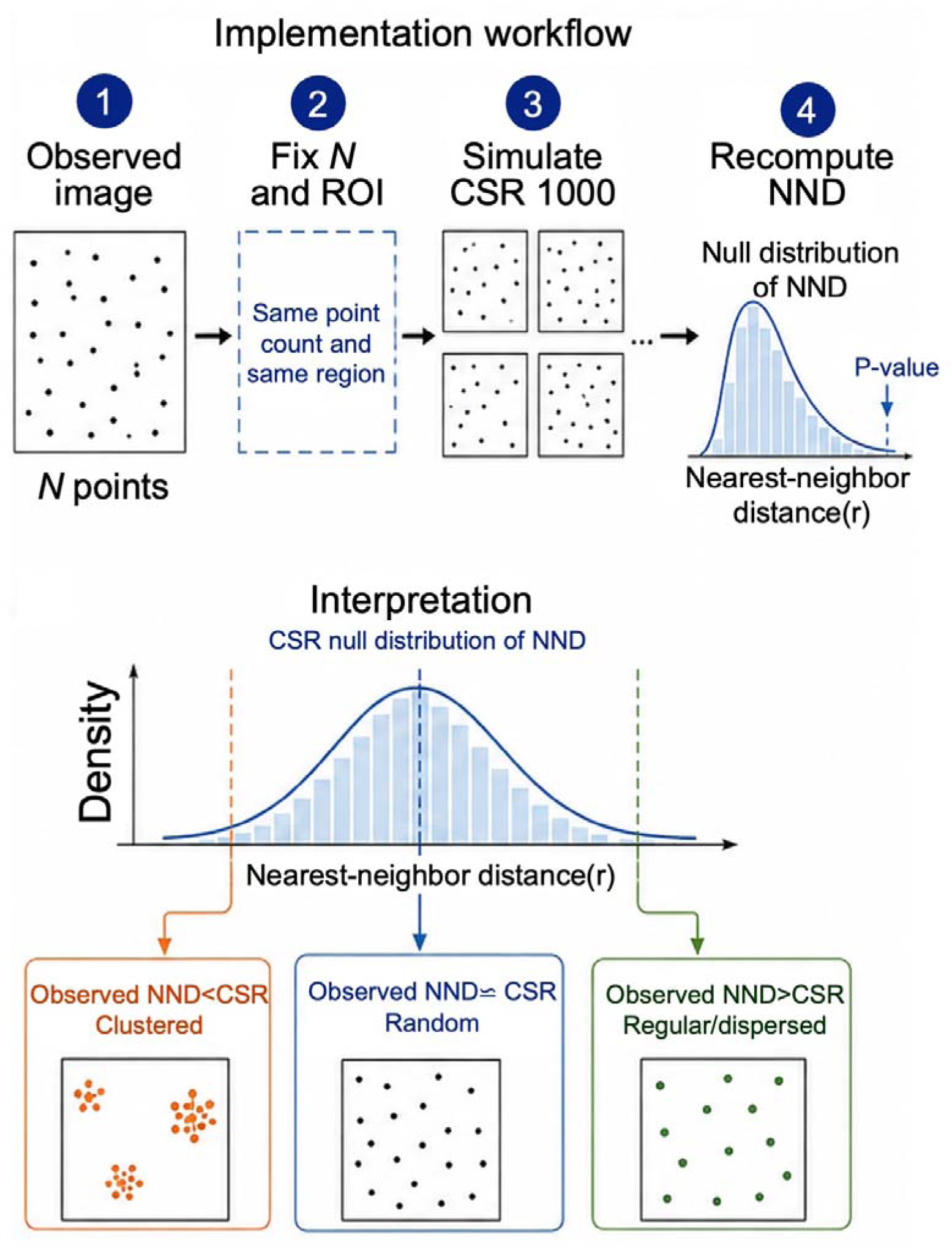
Complete spatial randomness null model used in the nearest-neighbor distance (NND) analysis of intracellular lipid droplets. Under complete spatial randomness (CSR), points (droplets) are distributed independently and uniformly over the intracellular observation window, with neither clustering nor repulsion. For each cell, N points are drawn uniformly within the same intracellular region, where N equals the observed droplet count, and the NND distribution is recomputed. The draw is repeated 1,000 times per cell to yield the CSR null distribution, whose median defines the CSR-expected median NND used in the per-cell R ratio (Methods, “Spatial point-pattern analysis of lipid droplets”). An observed median NND below the CSR envelope indicates clustering, one above it indicates regular or dispersed organization, and one within the envelope is indistinguishable from spatially random placement. The CSR null model and its implementation follow standard spatial point-pattern analysis^2^.

**Supplementary Fig. 7.**
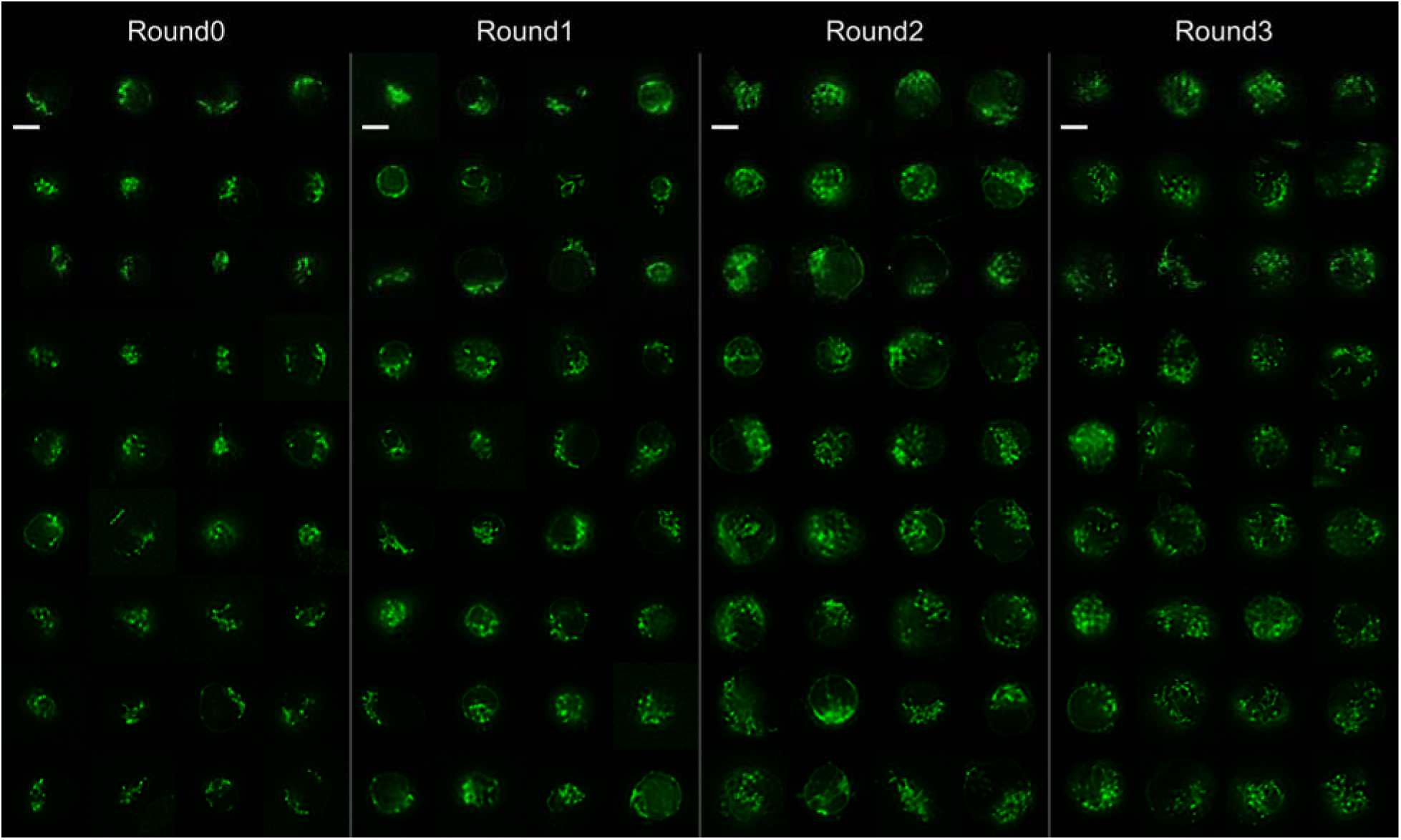
SR-IFC cell gallery for CD8^+^ T cells. Randomly sampled 16 by 9 montage of SR-IFC single-cell mitochondrial images across the four anti-CD3/CD28 stimulation rounds (48 h per round), showing the morphological heterogeneity of mitochondria. Read from round 0 to round 3, mitochondria shift from sparse, low-density distributions toward fragmented, punctate organelles, passing through a transient interconnected, network-like state. Scale bar, 5 μm.

**Supplementary Fig. 8.**
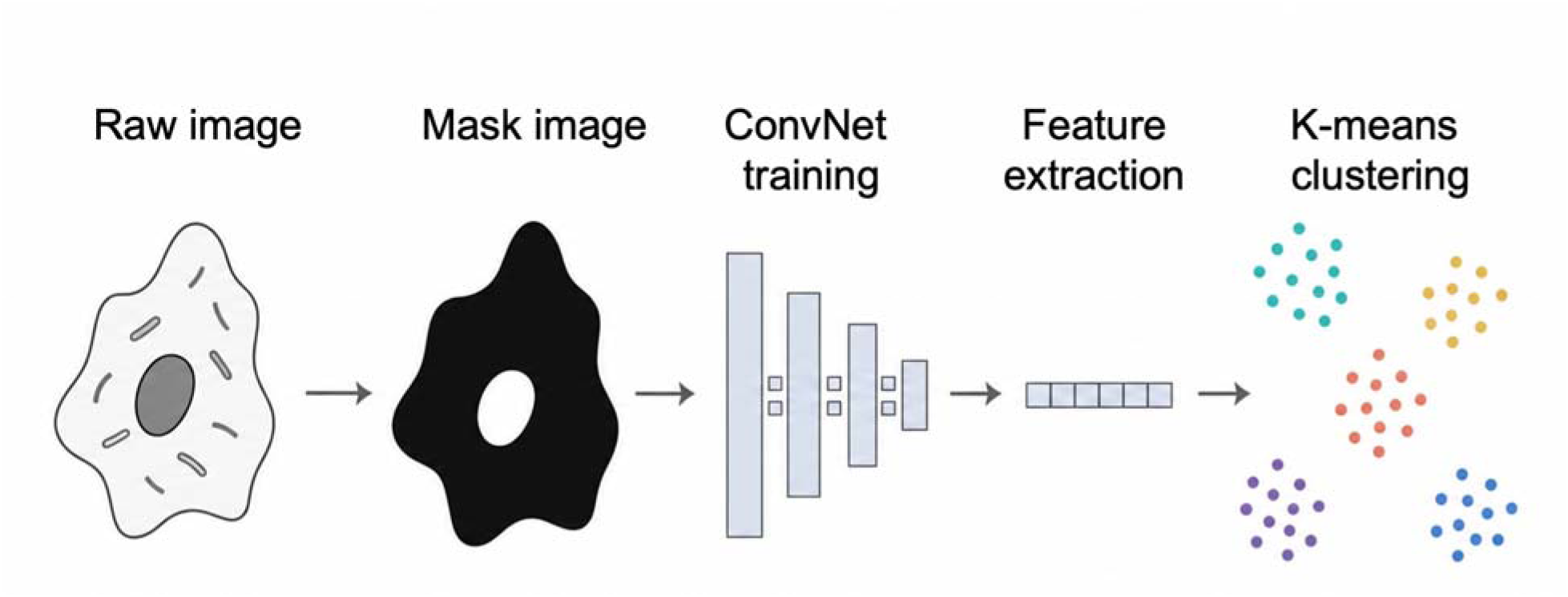
Self-supervised feature-clustering pipeline for mitochondrial morphology. Each SR-IFC single-cell image is restricted to a mitochondrial foreground mask generated by top-hat filtering, Otsu thresholding, and morphological cleanup (Methods, “Image segmentation and morphological analysis”). A ConvNeXt-Tiny backbone^3^ is trained on the masked images with a SimCLR^4^ self-supervised contrastive objective and produces one 768-dimensional embedding per cell. k-means clustering with *K* = 3 assigns the embeddings to the image-defined morphotypes C0 to C2. UMAP^5^ is used only for visualization (Fig. 5d).

**Supplementary Fig. 9.**
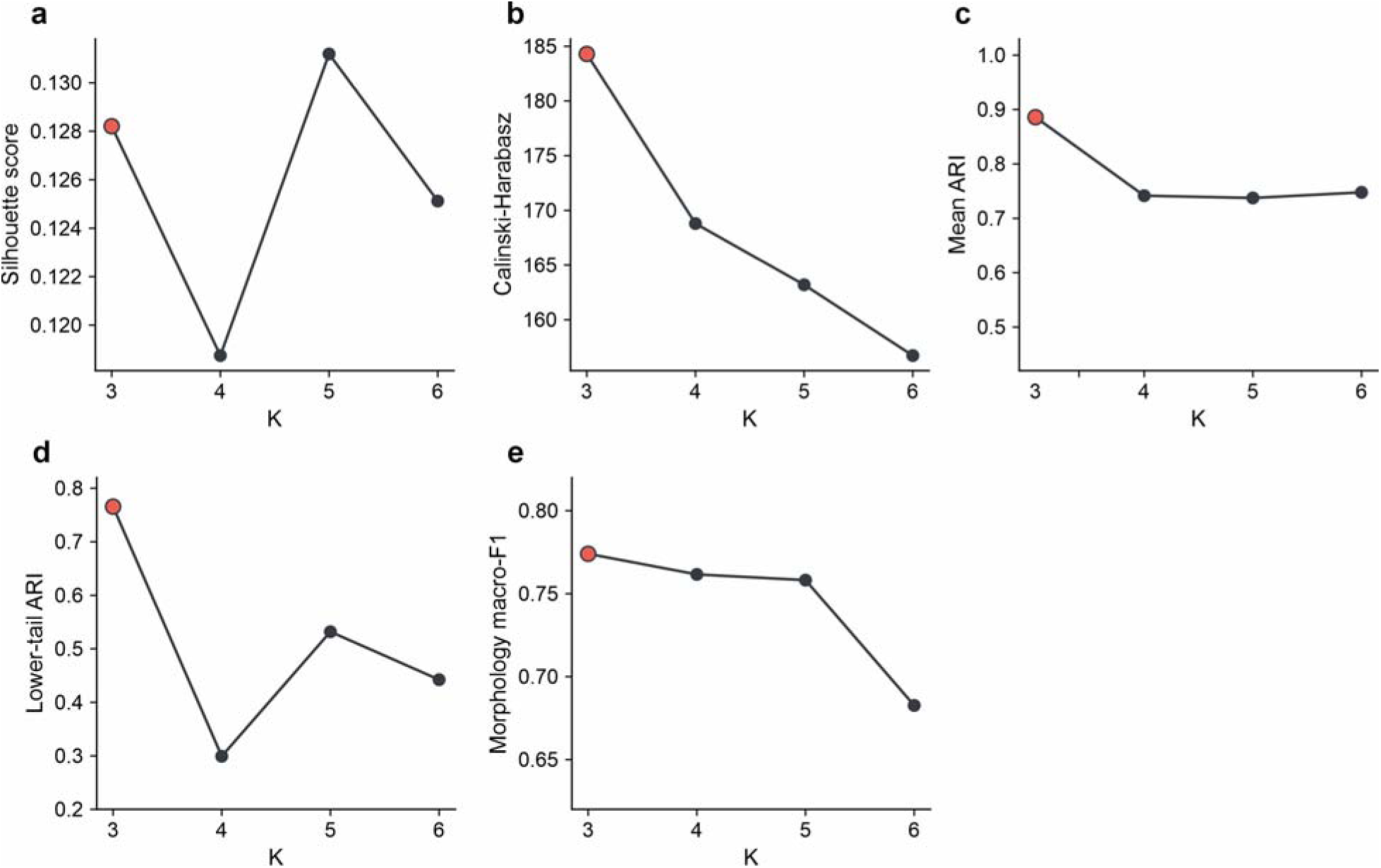
Data-driven selection of the number of mitochondrial morphotypes *K*. The number of morphotypes was determined from the data rather than imposed, using five criteria^6^ evaluated on the 768-dimensional embeddings of all SR-IFC single-cell images for *K* = 3 to 6. The adopted value, *K* = 3, is marked in red in every panel, and higher values are better throughout. **a**, Silhouette score^7^, the mean over cells of the normalized difference between each cell’s distance to its own group and to the nearest other group. **b**, Calinski-Harabasz score^8^, the ratio of between-group to within-group dispersion. **c**, Mean adjusted Rand index (ARI) between the full-data partition and 100 partitions obtained by refitting k-means on random 80% subsamples of the cells and reassigning all cells with the refitted centroids, measuring reproducibility under resampling of the cells. **d**, The 2.5th percentile of the same ARI distribution^9^, reporting reproducibility in the least favorable resamples; it is a fixed robustness metric, not a confidence interval. **e**, Mean macro-averaged F1 for predicting the morphotype labels from the eight morphometric features using class-balanced logistic regression in repeated stratified fivefold cross-validation (10 repeats, 50 held-out folds), measuring how well the morphotypes can be recovered from measurements that did not enter the clustering. *K* = 3 is highest in b to e. The silhouette score in a is marginally higher at *K* = 5 than at *K* = 3 (0.131 versus 0.128), a gain not accompanied by improved stability or recoverability. Silhouette values are low at every *K* tested (0.119 to 0.131), indicating a continuous morphology landscape rather than sharply separated classes; the three groups are therefore interpreted as reproducible morphotypes along that continuum.

**Supplementary Fig. 10.**
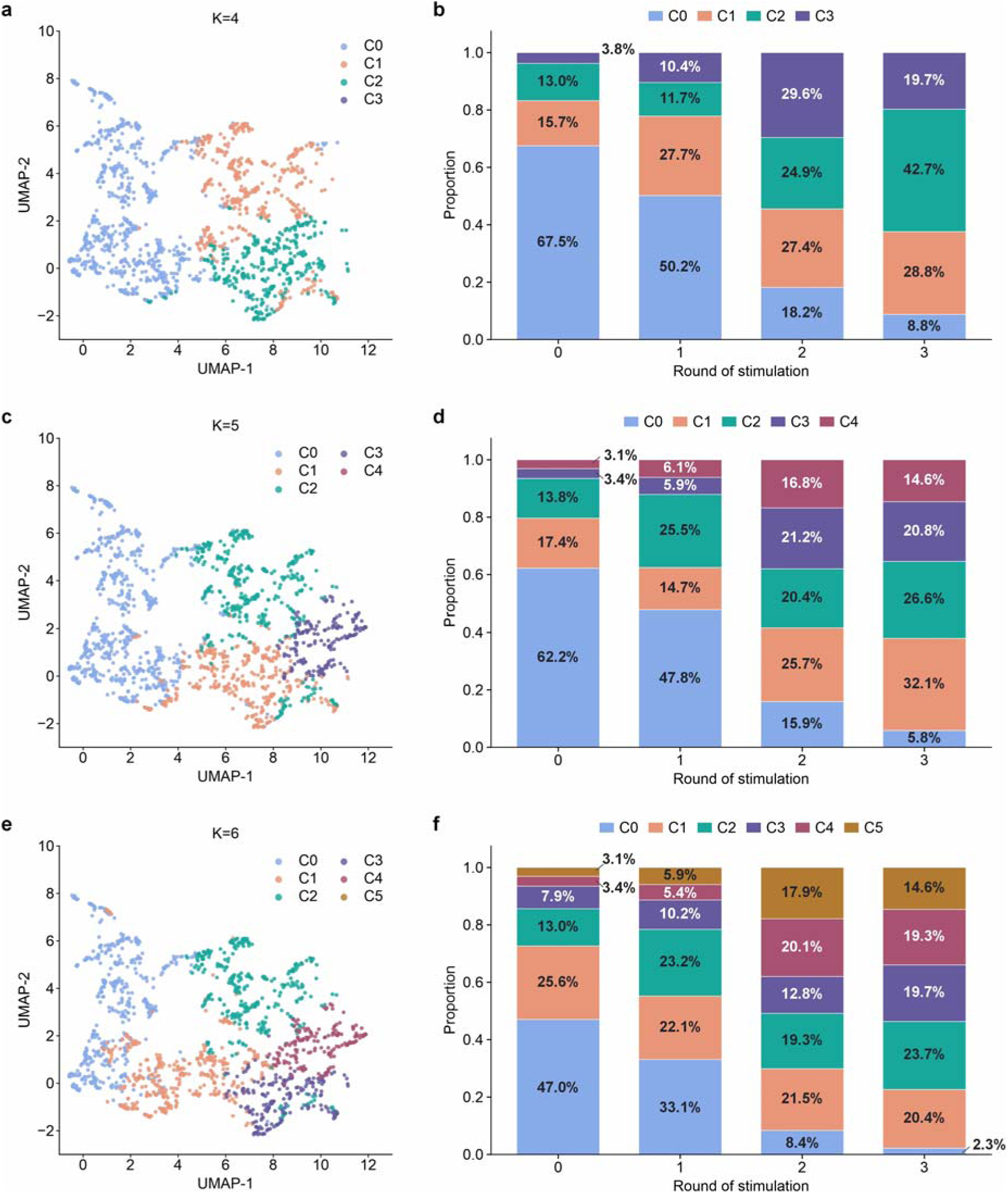
Mitochondrial-morphology clustering at *K* = 4 to 6. **a**, UMAP of the embeddings colored by the *K* = 4 clusters C0 to C3. **b**, Cluster composition after 0, 1, 2 and 3 rounds of stimulation for *K* = 4. **c**, UMAP colored by the *K* = 5 clusters C0 to C4. **d**, Cluster composition across rounds for *K* = 5. **e**, UMAP colored by the *K* = 6 clusters C0 to C5. **f**, Cluster composition across rounds for *K* = 6.

**Supplementary Fig. 11.**
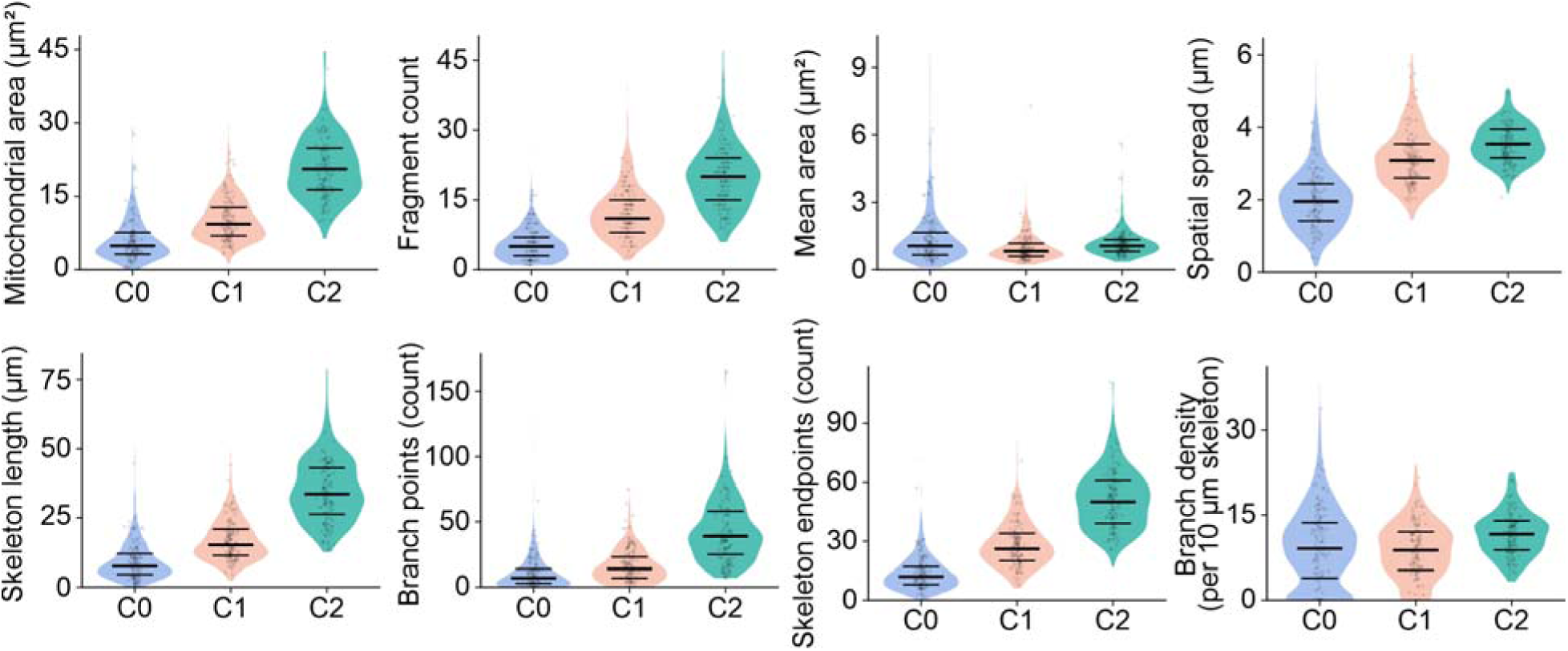
Per-feature morphometric characterization of the three mitochondrial morphotypes. To characterize the mitochondrial architectures associated with the self-supervised morphotypes C0 to C2, classical morphometric features were computed for the same cells from the same SR-IFC images, using the mitochondrial foreground segmentation described in Methods. Per-cell distributions across the three morphotypes are shown for eight features: total mitochondrial area, fragment count, mean fragment area, spatial spread, skeleton length, branch points, skeleton endpoints, and branch density. Violins show the per-cell distributions, horizontal lines mark the median and quartiles, and points show individual cells. Fig. 5e summarizes these features as morphotype-median z-scores standardized across C0 to C2, whereas this figure shows their full per-cell distributions.

**Supplementary Fig. 12.**
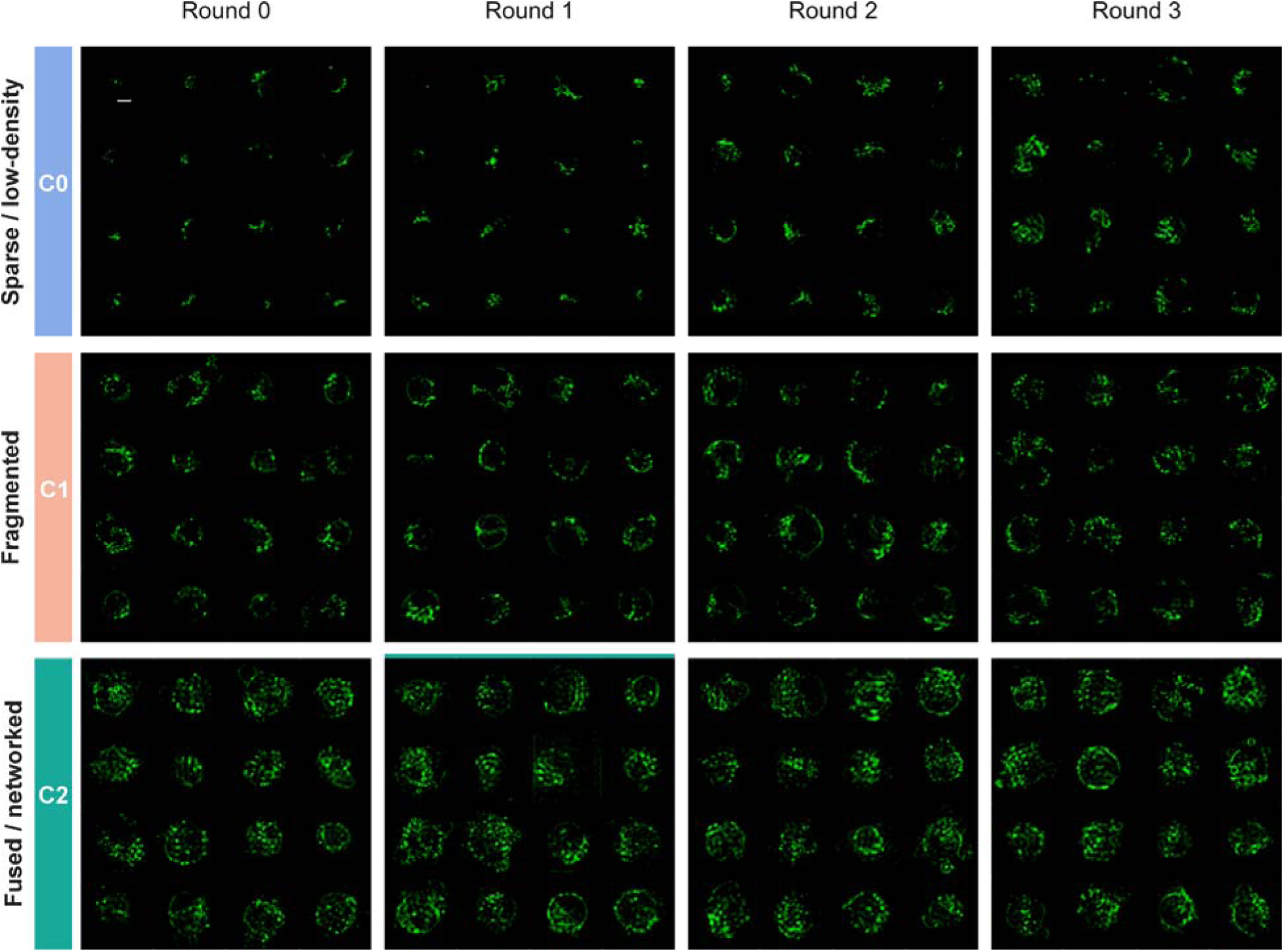
Per-morphotype mitochondrial image gallery across stimulation rounds. 4 × 4 grids of SR-IFC single-cell mitochondrial images sampled at random from each morphology morphotype, sparse/low-density (C0), fragmented (C1), and fused/networked (C2), after 0, 1, 2 and 3 rounds of anti-CD3/CD28 stimulation. Cells are sampled at random within each morphotype and round, giving an unbiased view of the typical morphology of each morphotype rather than hand-picked examples. Read down a column, the three morphotypes form the sparse-to-fragmented-to-fused gradient quantified in Fig. 5e and Supplementary Fig. 11; read across a row, the same morphotype is populated by different cells at every round, showing that morphotype assignment follows morphology rather than stimulation round. Scale bar, 5 μm.

**Supplementary Fig. 13.**
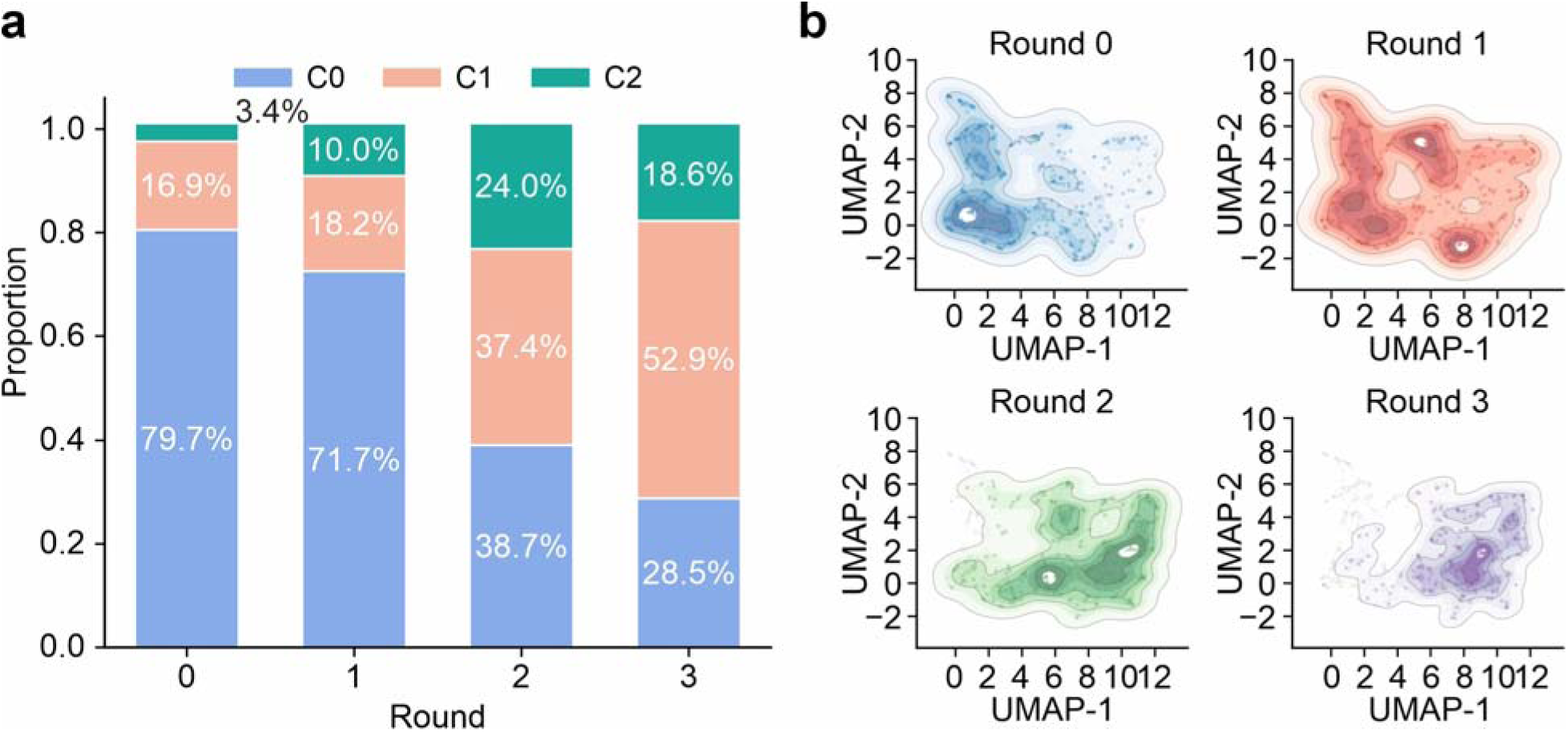
Round-resolved composition and UMAP occupancy of the three mitochondrial morphotypes at *K* = 3. **a**, Fraction of cells assigned to the sparse/low-density (C0), fragmented (C1), and fused/networked (C2) morphotypes after 0, 1, 2 and 3 rounds of stimulation. C0 falls from 79.7% to 28.5%, C2 rises to a transient peak of 24.0% at round 2 before declining to 18.6%, and C1 expands from 16.9% to 52.9% and becomes the dominant morphotype by round 3. **b**, Kernel density of the same cells in the UMAP embedding of Fig. 5d, shown separately for each round. The density shifts progressively over the three rounds without separating into discrete islands, consistent with a continuous morphology landscape.

**Supplementary Fig. 14.**
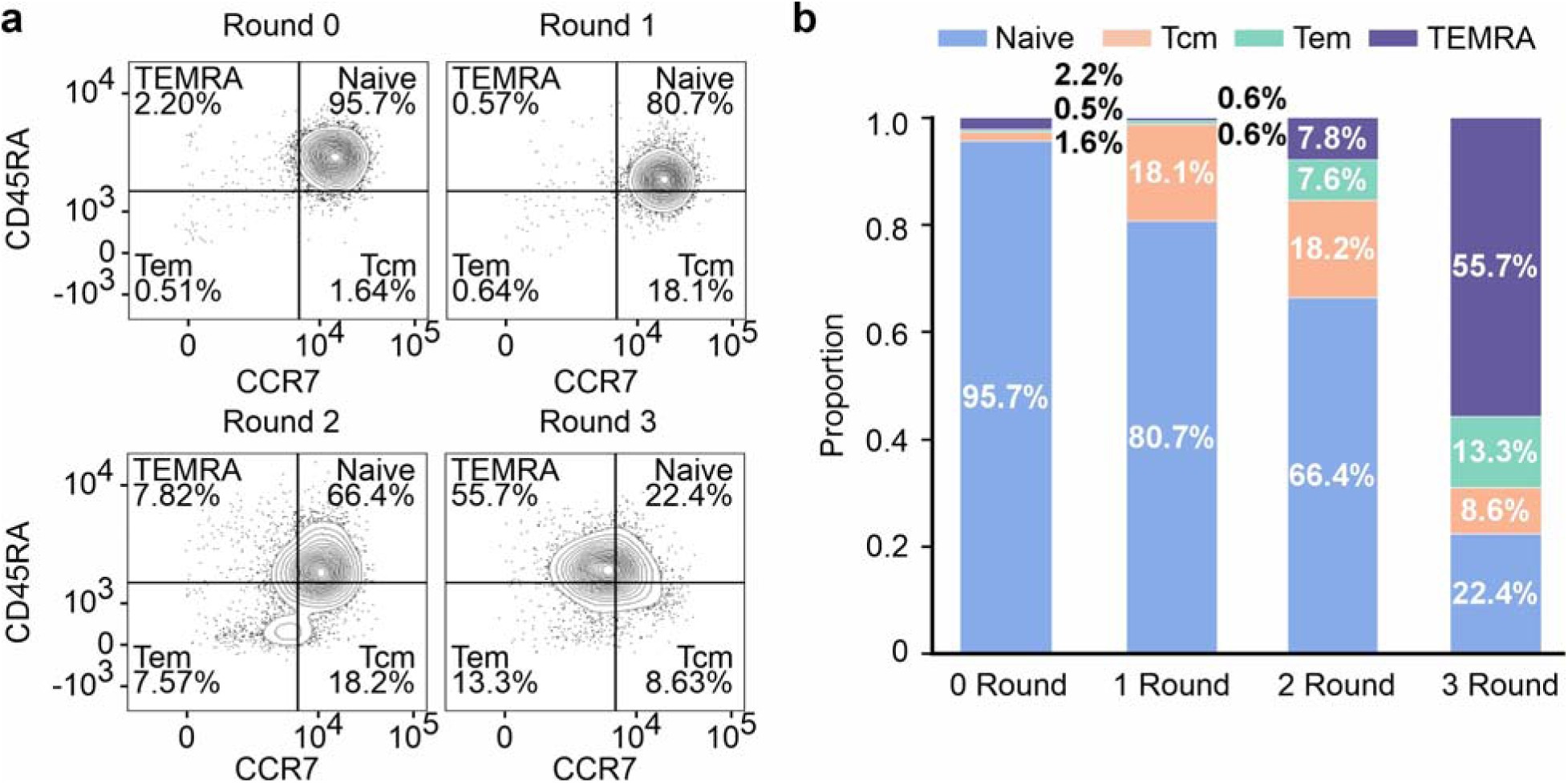
Matched flow cytometric analysis for Fig. 5. **a**, CD45RA versus CCR7 density plots of viable CD8^+^ T cells after 0, 1, 2, and 3 rounds of anti-CD3/CD28 stimulation. Quadrants define naive (CD45RA^+^CCR7^+^), central memory (Tcm, CD45RA^−^CCR7^+^), effector memory (Tem, CD45RA^−^CCR7^−^), and terminally differentiated effector memory cells re-expressing CD45RA (TEMRA, CD45RA^+^CCR7^−^) populations; percentages are shown within each quadrant. Quadrant thresholds were set with reference to CD45RA and CCR7 expression in naive T cells. **b**, Corresponding population proportions across the four stimulation rounds.

**Supplementary Fig. 15.**
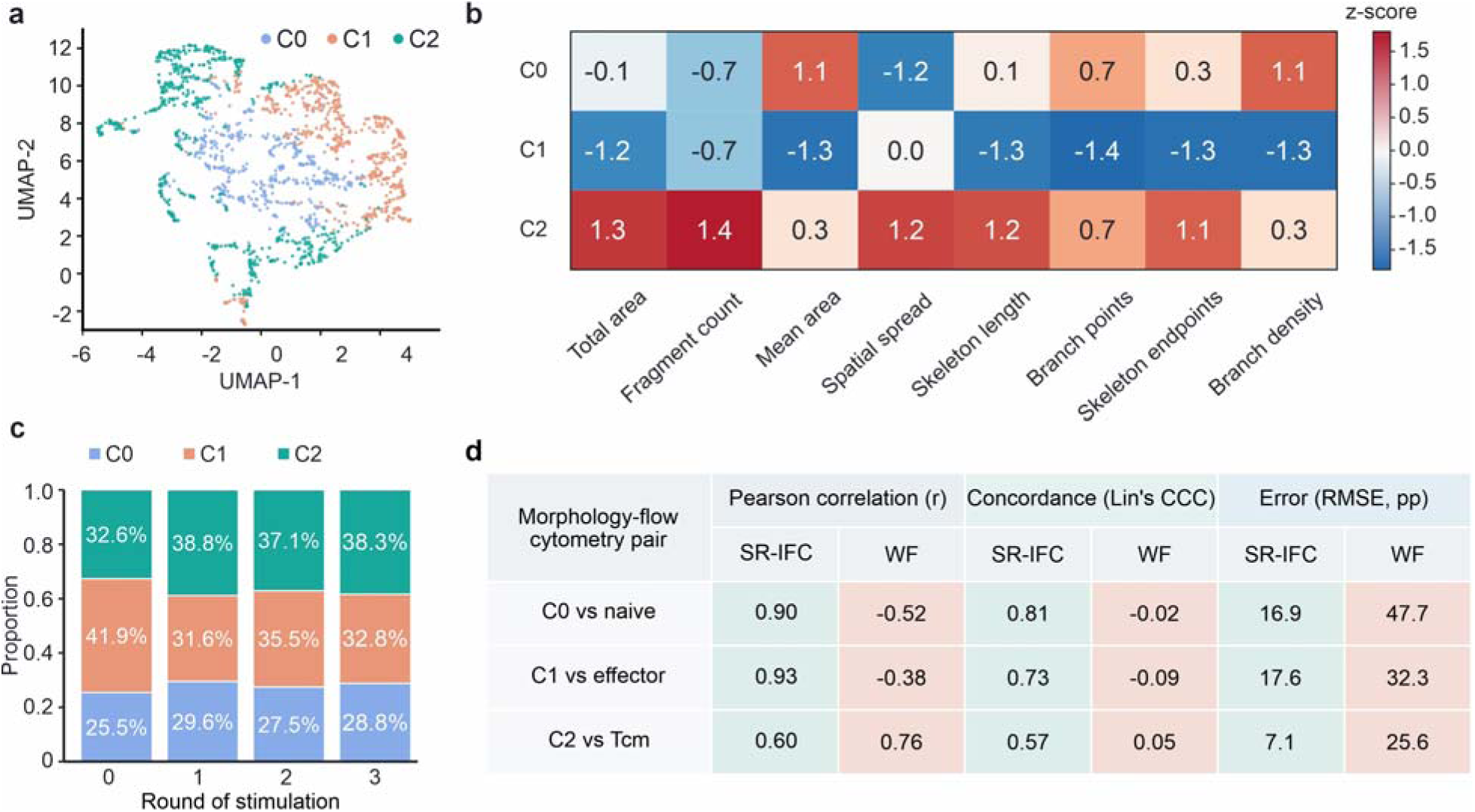
Effect of WF imaging on the inferred mitochondrial morphology trajectory. Matched WF images were analyzed with an independently trained WF-native encoder and the same downstream *K* = 3 clustering procedure; WF clusters were identified as the SR-IFC morphotypes. **a**, UMAP of the WF embeddings colored by the matched morphotype labels as the SR-IFC analysis. **b**, Morphotype-median z-scores for the eight morphometric features for C0 to C2. **c**, Proportions of C0 to C2 after 0, 1, 2, and 3 stimulation rounds. **d**, Pearson correlation coefficient (r), Lin’s concordance correlation coefficient (CCC), and root-mean-square error (RMSE, percentage points) for the prespecified pairs C0 versus naive, C1 versus effector (Tem and TEMRA pooled), and C2 versus Tcm, reported for SR-IFC and WF relative to the matched flow cytometric fractions.

**Supplementary Table 1.** Simulation parameters and noise model. Optical, noise, reconstruction, and ground-truth-geometry parameters used for the multi-scenario simulation (Fig. 2).

| **Optics and sampling** |  |
| --- | --- |
| **Parameter** | **Value** |
| emission wavelength | 525 nm |
| objective NA / refractive index | 1.40 / 1.515 |
| camera field | 256 × 256 pixels |
| camera pixel size | 65 nm |
| phantom pixel size | 16.25 nm |
| forward model | 3D incoherent projection: each z slice is convolved with a depth-dependent lateral PSF and then integrated at the camera |
| PSF model | Rayleigh-matched in-focus Gaussian |
| random seed | 20260505 |
| **Noise and image diversity** |  |
| **Parameter** | **Value** |
| noise model | Poisson photons + Gaussian read noise |
| read noise | 1.4 e<sup>-</sup> rms |
| bright-pixel signal | 180 e<sup>-</sup> |
| sample brightness | 0.60–1.50× |
| mode brightness | 0.75–1.25× |
| frame brightness | 0.90–1.10× |
| photon budget | 0.70–1.30× |
| Fig. 2e photon-noise sweep | Poisson only and peak relative shot noise from 0 to 50% |
| **SIM and reconstruction** |  |
| **Parameter** | **Value** |
| pattern period | 250 nm |
| modulation depth | 0.85 |
| SIM phases | 0, 2π/3, 4π/3 |
| flow-axis SIM orientation | 0° |
| conventional SIM orientations | 0, 60, 120° |
| Wiener / apodization | 0.08 / 100 nm |
| flow-axis SIM missing-direction rule | transverse-axis frequencies not covered by the single orientation are left at the WF support |

**Ground-truth geometries**
| Parameter | Value |
| --- | --- |
| bead pair (P1) | 5 × 5 grid of 25 pairs (50 spheres), each 100 nm in diameter; surface gaps = 250, 180, 120, 90 and 60 nm; orientations = 0°, 22.5°, 45°, 67.5° and 90°; centered at z = 0 with finite 3D extent |
| ellipsoid (P2) | 20 finite 3D ellipsoids; lateral aspect ratios = 1:1, 1:3, 1:5 and 1:10; orientations = 0°, 30°, 45°, 60° and 90°; minor-axis diameter = 90 to 110 nm (nominally 100 nm); centered at z = 0 |
| frequency target (P3) | 16 finite-thickness periodic line-pattern blocks; periods = 240, 300, 400 and 600 nm ($\pm 2\%$); orientations = 0°, 45°, 90° and 135° ($\pm 2^\circ$); block width = 1.65 to 1.95 $\mu\text{m}$; slab thickness = 130 to 180 nm; centered at z = 0 |

## Supplementary Note 1 Two-stage rigid registration of consecutive flowing-cell frames

For each cell, three phase-shifted frames *I□*(*r*), n = 1, 2, 3, are acquired stroboscopically at 720 frames per second. Within the short inter-frame interval, the cell moves to a different position in the camera ROI and may also undergo a small angular drift. Before SIM reconstruction, SR-IFC therefore corrects this rigid inter-frame motion in two stages, translation and then rotation, both based on Fourier cross-correlation^10^. Supplementary Fig. 3 illustrates the translation stage (Stage 1), from two consecutive flowing-cell frames *I□* and *I□□□*, through their Cartesian cross-correlation map C(δ), to the magnified correlation peak and the local fit that recovers the subpixel inter-frame translation.

Stage 1: translation correction. Because the cell occupies different positions in the camera ROI in consecutive frames, the Cartesian Fourier cross-correlation between *I□* and *I□□□* is computed as

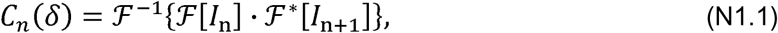

where *F* is the two-dimensional Fourier transform and * denotes complex conjugation. The integer-pixel displacement is the global maximum,

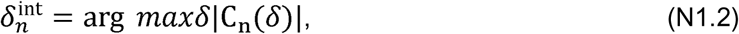

with subpixel refinement obtained by a two-dimensional Gaussian (log-quadratic) fit over the 3 × 3 neighborhood of 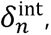

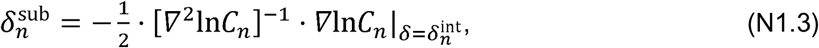

yielding the full translation 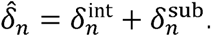. *I□□□* is then shifted by − *δ̂_n_* using spline interpolation to bring it into the reference frame of *I□*.

Stage 2: rotation correction^11^. The 720 frames per second frame rate keeps the inter-frame angular drift small (typically below 0.5°). To recover this sub-degree rotation reliably, the translation-aligned pair is first upsampled twofold in both dimensions by bicubic interpolation. The Fourier magnitudes of the upsampled frames are then remapped to log-polar coordinates (ρ, θ) = (log |*k*|,□*k*), in which an in-plane rotation becomes a pure translation along the θ axis (Fourier–Mellin construction),

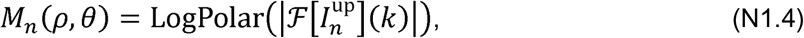

where 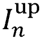 denotes the twofold upsampled frame. Cross-correlating *M□* and *M□□□* along θ and locating the maximum yields the inter-frame rotation angle,

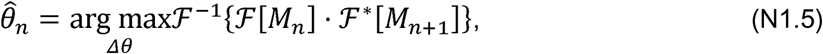

and the upsampled, translation-aligned *I□□□* is rotated by – *θ̂_n_* about its centroid before being downsampled to the native pixel grid. The combined rigid transform (translation by − *δ̂_n_* followed by rotation by − *θ̂_n_* in the upsampled domain) registers all three phase frames to the reference frame of *I□*. The registered triplet is then averaged to form the WF image and passed to the blind-SIM reconstruction of Supplementary Note 2 to form the flow-axis SIM image.

## Supplementary Note 2 Single-orientation blind-SIM reconstruction along the modulation axis

SR-IFC reconstructs the flow-axis super-resolved image (flow-axis SIM) from three registered fluorescence frames acquired with a single sinusoidal illumination orientation. Standard three-phase SIM assumes nominal phase offsets of 0°, 120°, and 240°^12–14^. During continuous-flow acquisition, residual errors in PZT phase stepping, acquisition timing, and frame registration can make the effective phase offsets cell dependent. We therefore use a parametric blind-SIM formulation in which the fluorophore distribution and illumination parameters are jointly estimated^15^. Let *r* = (*r*___, *r*___) denote sample-plane coordinates, with *r* parallel to the modulation (flow) axis. The *j*th registered frame is modeled as

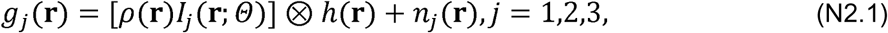

where ρ is the non-negative fluorophore distribution, *h* is the point-spread function (PSF), *n*_j_ is measurement noise, and Θ denotes the illumination parameters. The sinusoidal illumination and effective phase of each frame are

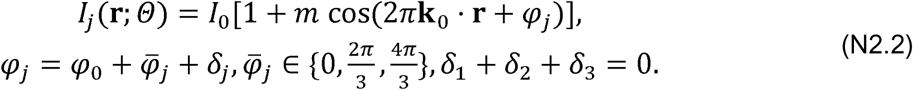

Here Θ = (*I*_0_, *m*, *k*_0_, φ_0_, δ_1_, δ_2_, δ_3_) comprises the mean intensity, modulation depth, carrier vector, common phase, and frame-specific phase-step deviations. The zero-sum constraint separates the common phase from the relative phase errors. Fourier transformation of Eq. (N2.1) gives

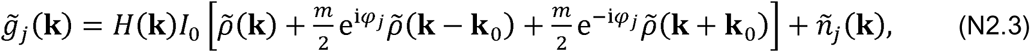

which is a 3 × 3 mixing relation among the central and ±1 orders. Unlike conventional SIM, the phase matrix is determined by the jointly estimated Θ rather than by the nominal phases alone. Sample positivity is enforced through ρ = ξ*^2^*. With □*_j_* denoting multiplication by the current illumination followed by PSF convolution, the joint estimate is obtained from

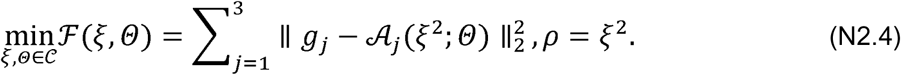

The feasible set□ enforces 0 < *m* ≤ 1, the zero-sum phase constraint, and carrier frequency and orientation within the calibrated neighborhood. Each iteration applies two Polak-Ribière conjugate-gradient sweeps, one over the sample variable and one over the illumination parameters, with line-search step sizes:

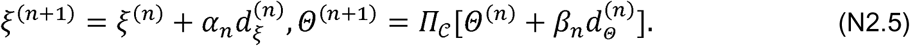

The Polak-Ribière directions are formed from the corresponding gradients of □, and the projection Π□ restores the illumination constraints. The reconstruction typically converges within approximately 15 iterations to a normalized data residual below 10^−3^. After convergence, the estimated phase matrix separates the central and ±1 orders; these components are Wiener-filtered with the calibrated optical transfer function (OTF), shifted to their true frequency positions, and recombined to form the flow-axis SIM image. This step is a conventional SIM frequency-component recombination and involves no network output. It extends the OTF support along the modulation (flow) axis while leaving the orthogonal axis diffraction limited. The registered WF image and the resulting flow-axis SIM image are then supplied as the two inputs to the dual-input network described in Methods (“AI-assisted isotropic reconstruction”) to generate the final SR-IFC image with isotropic in-plane resolution.

## REFERENCES

1. Raj, A. & van Oudenaarden, A. Nature, nurture, or chance: stochastic gene expression and its consequences. Cell 135, 216–226 (2008).

2. Luzak, V., López-Escobar, L., Siegel, T. N. & Figueiredo, L. M. Cell-to-cell heterogeneity in trypanosomes. Annu. Rev. Microbiol. 75, 107–128 (2021).

3. Cohen, S., Valm, A. M. & Lippincott-Schwartz, J. Interacting organelles. Curr. Opin. Cell Biol. 53, 84–91 (2018).

4. Chang, A. Y. & Marshall, W. F. Organelles – understanding noise and heterogeneity in cell biology at an intermediate scale. J. Cell Sci. 130, 819–826 (2017).

5. Rafelski, S. M. & Marshall, W. F. Building the cell: design principles of cellular architecture. Nat. Rev. Mol. Cell Biol. 9, 593–602 (2008).

6. Valm, A. M. et al. Applying systems-level spectral imaging and analysis to reveal the organelle interactome. Nature 546, 162–167 (2017).

7. Altschuler, S. J. & Wu, L. F. Cellular heterogeneity: do differences make a difference? Cell 141, 559–563 (2010).

8. Herbig, M. et al. Best practices for reporting throughput in biomedical research. Nat. Methods 19, 633–634 (2022).

9. Sahl, S. J., Hell, S. W. & Jakobs, S. Fluorescence nanoscopy in cell biology. Nat. Rev. Mol. Cell Biol. 18, 685–701 (2017).

10. Barentine, A. E. S. et al. An integrated platform for high-throughput nanoscopy. Nat. Biotechnol. 41, 1549–1556 (2023).

11. Beghin, A. et al. Localization-based super-resolution imaging meets high-content screening. Nat. Methods 14, 1184–1190 (2017).

12. Guo, M. et al. High-speed super-resolution structured illumination microscopy with a large field-of-view. Opt. Express 32, 31485–31494 (2024).

13. Burns, Z., Zhao, J., Sahan, A. Z., Zhang, J. & Liu, Z. High-speed blind structured illumination microscopy via unsupervised algorithm unrolling. Nat. Commun. 17, 1967 (2026).

14. Zhao, W. et al. Enhanced detection of fluorescence fluctuations for high-throughput super-resolution imaging. Nat. Photon. 17, 806–813 (2023).

15. Yoon, K. et al. Real-time, high-throughput super-resolution microscopy via panoramic integration. Nat. Commun. 16, 9309 (2025).

16. Han, Y., Gu, Y., Zhang, A. C. & Lo, Y.-H. Review: imaging technologies for flow cytometry. Lab Chip 16, 4639–4647 (2016).

17. Goda, K., et al. High-throughput single-microparticle imaging flow analyzer. Proc. Natl Acad. Sci. USA 109, 11630–11635 (2012).

18. Lei, C. et al. High-throughput imaging flow cytometry by optofluidic time-stretch microscopy. Nat. Protoc. 13, 1603–1631 (2018).

19. Lei, C. et al. High-throughput optical imaging technology for large-scale single-cell analysis of live *Euglena gracilis*. TrAC Trends Anal. Chem. 180, 117938 (2024).

20. Rees, P., Summers, H. D., Filby, A., Carpenter, A. E. & Doan, M. Imaging flow cytometry. Nat. Rev. Methods Primers 2, 86 (2022).

21. Suzuki, Y., et al. Label-free chemical imaging flow cytometry by high-speed multicolor stimulated Raman scattering. Proc. Natl Acad. Sci. USA 116, 15842–15848 (2019).

22. Miura, T. et al. On-chip light-sheet fluorescence imaging flow cytometry at a high flow speed of 1 m/s. Biomed. Opt. Express 9, 3424–3433 (2018).

23. Kanno, H. et al. High-throughput fluorescence lifetime imaging flow cytometry. Nat. Commun. 15, 7376 (2024).

24. Hua, X. et al. Light-field flow cytometry for high-resolution, volumetric and multiparametric 3D single-cell analysis. Nat. Commun. 15, 1975 (2024).

25. Holzner, G. et al. High-throughput multiparametric imaging flow cytometry: toward diffraction-limited sub-cellular detection and monitoring of sub-cellular processes. Cell Rep. 34, 108824 (2021).

26. Weiss, L. E. et al. Three-dimensional localization microscopy in live flowing cells. Nat. Nanotechnol. 15, 500–506 (2020).

27. Mandracchia, B., Son, J. & Jia, S. Super-resolution optofluidic scanning microscopy. Lab Chip 21, 489–493 (2021).

28. Paiè, P. et al. Structured-light-sheet imaging in an integrated optofluidic platform. Lab Chip 24, 34–46 (2024).

29. Sigal, Y. M., Zhou, R. & Zhuang, X. Visualizing and discovering cellular structures with super-resolution microscopy. Science 361, 880–887 (2018).

30. Betzig, E. et al. Imaging intracellular fluorescent proteins at nanometer resolution. Science 313, 1642–1645 (2006).

31. Rust, M. J., Bates, M. & Zhuang, X. Sub-diffraction-limit imaging by stochastic optical reconstruction microscopy (STORM). Nat. Methods 3, 793–796 (2006).

32. Hell, S. W. & Wichmann, J. Breaking the diffraction resolution limit by stimulated emission: stimulated-emission-depletion fluorescence microscopy. Opt. Lett. 19, 780–782 (1994).

33. Hell, S. W. Far-field optical nanoscopy. Science 316, 1153–1158 (2007).

34. Gustafsson, M. G. L. Surpassing the lateral resolution limit by a factor of two using structured illumination microscopy. J. Microsc. 198, 82–87 (2000).

35. Schermelleh, L., et al. Super-resolution microscopy demystified. Nat. Cell Biol. 21, 72–84 (2019).

36. Mao, X., Lin, S.-C. S., Dong, C. & Huang, T. J. Single-layer planar on-chip flow cytometer using microfluidic drifting based three-dimensional (3D) hydrodynamic focusing. Lab Chip 9, 1583– 1589 (2009).

37. Gustafsson, M. G. L. et al. Three-dimensional resolution doubling in wide-field fluorescence microscopy by structured illumination. Biophys. J. 94, 4957–4970 (2008).

38. Lal, A., Shan, C. & Xi, P. Structured illumination microscopy image reconstruction algorithm. IEEE J. Sel. Top. Quantum Electron. 22, 50–63 (2016).

39. Ayuk, R. et al. Structured illumination fluorescence microscopy with distorted excitations using a filtered blind-SIM algorithm. Opt. Lett. 38, 4723–4726 (2013).

40. Wang, H. et al. Deep learning enables cross-modality super-resolution in fluorescence microscopy. Nat. Methods 16, 103–110 (2019).

41. Belthangady, C. & Royer, L. A. Applications, promises, and pitfalls of deep learning for fluorescence image reconstruction. Nat. Methods 16, 1215–1225 (2019).

42. Ronneberger, O., Fischer, P. & Brox, T. U-Net: convolutional networks for biomedical image segmentation. In Medical Image Computing and Computer-Assisted Intervention – MICCAI 2015 (eds Navab, N., Hornegger, J., Wells, W. M. & Frangi, A. F.) Vol. 9351, 234–241 (Springer, 2015).

43. Qiao, C. et al. Evaluation and development of deep neural networks for image super-resolution in optical microscopy. Nat. Methods 18, 194–202 (2021).

44. Wang, Z., Simoncelli, E. P. & Bovik, A. C. Multiscale structural similarity for image quality assessment. In Proc. 37th Asilomar Conference on Signals, Systems and Computers Vol. 2, 1398–1402 (IEEE, 2003).

45. Kim, T. H. & Haldar, J. P. The Fourier radial error spectrum plot: a more nuanced quantitative evaluation of image reconstruction quality. In 2018 IEEE 15th International Symposium on Biomedical Imaging (ISBI 2018) 61–64 (IEEE, 2018).

46. Jin, L. et al. Deep learning enables structured illumination microscopy with low light levels and enhanced speed. Nat. Commun. 11, 1934 (2020).

47. Thiam, A. R., Farese, R. V. & Walther, T. C. The biophysics and cell biology of lipid droplets. Nat. Rev. Mol. Cell Biol. 14, 775–786 (2013).

48. Goold, H., Beisson, F., Peltier, G. & Li-Beisson, Y. Microalgal lipid droplets: composition, diversity, biogenesis and functions. Plant Cell Rep. 34, 545–555 (2015).

49. Sun, X.-M., Ren, L.-J., Zhao, Q.-Y., Ji, X.-J. & Huang, H. Microalgae for the production of lipid and carotenoids: a review with focus on stress regulation and adaptation. Biotechnol. Biofuels 11, 272 (2018).

50. Shitanaka, T. et al. Synthetic biology and metabolic engineering of microalgae for sustainable lipid and terpenoid production: an updated perspective. Plant Biotechnol. J. 24, 1359–1376 (2026).

51. Baddeley, A. & Turner, R. spatstat: an R package for analyzing spatial point patterns. J. Stat. Softw. 12, 1–42 (2005).

52. Bohnert, M. Tether me, tether me not—dynamic organelle contact sites in metabolic rewiring. Dev. Cell 54, 212–225 (2020).

53. Walther, T. C., Chung, J. & Farese, R. V. Lipid droplet biogenesis. Annu. Rev. Cell Dev. Biol. 33, 491–510 (2017).

54. Buck, M. D. et al. Mitochondrial dynamics controls T cell fate through metabolic programming. Cell 166, 63–76 (2016).

55. van der Windt, G. J. W. et al. Mitochondrial respiratory capacity is a critical regulator of CD8^+^ T cell memory development. Immunity 36, 68–78 (2012).

56. O’Sullivan, D. et al. Memory CD8^+^ T cells use cell-intrinsic lipolysis to support the metabolic programming necessary for development. Immunity 41, 75–88 (2014).

57. Mishra, P. & Chan, D. C. Metabolic regulation of mitochondrial dynamics. J. Cell Biol. 212, 379– 387 (2016).

58. Fischer, M. et al. Early effector maturation of naïve human CD8^+^ T cells requires mitochondrial biogenesis. Eur. J. Immunol. 48, 1632–1643 (2018).

59. Klein Geltink, R. I., et al. Mitochondrial priming by CD28. Cell 171, 385–397.e11 (2017).

60. Trickett, A. & Kwan, Y. L. T cell stimulation and expansion using anti-CD3/CD28 beads. J. Immunol. Methods 275, 251–255 (2003).

61. Rambold, A. S., Kostelecky, B., Elia, N. & Lippincott-Schwartz, J. Tubular network formation protects mitochondria from autophagosomal degradation during nutrient starvation. Proc. Natl Acad. Sci. USA 108, 10190–10195 (2011).

62. Liu, Z. et al. A ConvNet for the 2020s. In 2022 IEEE/CVF Conference on Computer Vision and Pattern Recognition (CVPR) 11966–11976 (IEEE, 2022).

63. Chen, T., Kornblith, S., Norouzi, M. & Hinton, G. A simple framework for contrastive learning of visual representations. In Proc. 37th International Conference on Machine Learning (eds Daumé III, H. & Singh, A.) Vol. 119, 1597–1607 (PMLR, 2020).

64. Sallusto, F., Lenig, D., Förster, R., Lipp, M. & Lanzavecchia, A. Two subsets of memory T lymphocytes with distinct homing potentials and effector functions. Nature 401, 708–712 (1999).

65. Hamann, D. et al. Phenotypic and functional separation of memory and effector human CD8^+^ T cells. J. Exp. Med. 186, 1407–1418 (1997).

66. Poon, M. M. L. et al. Tissue adaptation and clonal segregation of human memory T cells in barrier sites. Nat. Immunol. 24, 309–319 (2023).

67. Beura, L. K. et al. Normalizing the environment recapitulates adult human immune traits in laboratory mice. Nature 532, 512–516 (2016).

68. Simula, L. et al. Drp1 controls effective T cell immune-surveillance by regulating T cell migration, proliferation, and cMyc-dependent metabolic reprogramming. Cell Rep. 25, 3059–3073.e10 (2018).

69. Quintana, A., et al. T cell activation requires mitochondrial translocation to the immunological synapse. Proc. Natl Acad. Sci. USA 104, 14418–14423 (2007).

70. Humblin, E., et al. Sustained CD28 costimulation is required for self-renewal and differentiation of TCF-1^+^ PD-1^+^ CD8 T cells. Sci. Immunol. 8, eadg0878 (2023).

71. Yu, Y.-R. et al. Disturbed mitochondrial dynamics in CD8^+^ TILs reinforce T cell exhaustion. Nat. Immunol. 21, 1540–1551 (2020).

72. Nitta, N. et al. Intelligent image-activated cell sorting. Cell 175, 266–276.e13 (2018).

73. Ding, T., et al. Image-activated cell sorting. Nat. Rev. Bioeng. 3, 890–907 (2025).

## REFERENCES

1. Ronneberger, O., Fischer, P. & Brox, T. U-Net: convolutional networks for biomedical image segmentation. In Medical Image Computing and Computer-Assisted Intervention – MICCAI 2015 (eds Navab, N., Hornegger, J., Wells, W. M. & Frangi, A. F.) Vol. 9351, 234–241 (Springer, 2015).

2. Baddeley, A. & Turner, R. spatstat: an R package for analyzing spatial point patterns. J. Stat. Softw. 12, 1–42 (2005).

3. Liu, Z. et al. A ConvNet for the 2020s. In 2022 IEEE/CVF Conference on Computer Vision and Pattern Recognition (CVPR) 11966–11976 (IEEE, 2022).

4. Chen, T., Kornblith, S., Norouzi, M. & Hinton, G. A simple framework for contrastive learning of visual representations. In Proc. 37th International Conference on Machine Learning (eds Daumé III, H. & Singh, A.) Vol. 119, 1597–1607 (PMLR, 2020).

5. Healy, J. & McInnes, L. Uniform manifold approximation and projection. Nat. Rev. Methods Primers 4, 82 (2024).

6. Ashari, I. F., Dwi Nugroho, E., Baraku, R., Novri Yanda, I. & Liwardana, R. Analysis of elbow, silhouette, Davies-Bouldin, Calinski-Harabasz, and Rand-index evaluation on k-means algorithm for classifying flood-affected areas in Jakarta. J. Appl. Inform. Comput. 7, 95–103 (2023).

7. Rousseeuw, P. J. Silhouettes: a graphical aid to the interpretation and validation of cluster analysis. J. Comput. Appl. Math. 20, 53–65 (1987).

8. Caliński, T. & Harabasz, J. A dendrite method for cluster analysis. Commun. Stat. 3, 1–27 (1974).

9. Rand, W. M. Objective criteria for the evaluation of clustering methods. J. Am. Stat. Assoc. 66, 846–850 (1971).

10. Reddy, B. S. & Chatterji, B. N. An FFT-based technique for translation, rotation, and scale-invariant image registration. IEEE Trans. Image Process. 5, 1266–1271 (1996).

11. Tzimiropoulos, G., Argyriou, V., Zafeiriou, S. & Stathaki, T. Robust FFT-based scale-invariant image registration with image gradients. IEEE Trans. Pattern Anal. Mach. Intell. 32, 1899–1906 (2010).

12. Gustafsson, M. G. L. Surpassing the lateral resolution limit by a factor of two using structured illumination microscopy. J. Microsc. 198, 82–87 (2000).

13. Gustafsson, M. G. L. et al. Three-dimensional resolution doubling in wide-field fluorescence microscopy by structured illumination. Biophys. J. 94, 4957–4970 (2008).

14. Lal, A., Shan, C. & Xi, P. Structured illumination microscopy image reconstruction algorithm. IEEE J. Sel. Top. Quantum Electron. 22, 50–63 (2016).

15. Ayuk, R. et al. Structured illumination fluorescence microscopy with distorted excitations using a filtered blind-SIM algorithm. Opt. Lett. 38, 4723–4726 (2013).

